# Small Molecule Targeting IRES Domain Inhibits Enterovirus 71 Replication via an Allosteric Mechanism that Stabilizes a Ternary Complex

**DOI:** 10.1101/2020.03.10.981167

**Authors:** Jesse Davila-Calderon, Neeraj Patwardhan, Liang-Yuan Chiu, Andrew Sugarman, Zhengguo Cai, Srinivasa R. Penutmutchu, Mei-Ling Li, Gary Brewer, Amanda E. Hargrove, Blanton S. Tolbert

## Abstract

We herein report an RNA-targeting antiviral small molecule that reduces replication of the human enterovirus 71 (EV71) via stabilization of an inhibitory small molecule-RNA-protein ternary complex. The EV71 virus poses serious threats to human health, particularly in regions of Southeast Asia, and no FDA approved drugs or vaccines are available. We first screened an RNA-biased small molecule library using a peptide-displacement assay to identify ligands for the stem loop II structure of the EV71 internal ribosomal entry site, which was previously shown to impact viral translation and replication. One ligand, DMA-135, decreased viral translation and replication in cell-based studies in a dose-dependent manner with no significant toxicity. Structural, biophysical, and biochemical characterization support an allosteric mechanism in which DMA-135 induces a conformational change in the RNA structure that stabilizes a ternary complex with the AUF1 protein that then represses translation. This mechanism was further supported by pull-down experiments in cell culture. These detailed studies establish enterovirus RNA structures as promising drug targets while revealing an approach and mechanism of action that should be broadly applicable to functional RNA targeting.

## Introduction

Human enterovirus 71 (EV71) is one of the major etiological agents of hand, foot, and mouth disease among children worldwide. Typical EV71 infections manifest with mild flu-like symptoms that can be overcome by the immune system; however, more severe cases can lead to neurological disorders, paralysis, respiratory failure and even death. EV71 outbreaks in parts of the Asia-Pacific, such as a recent (2018) epidemic in Vietnam where 53,000 children were hospitalized and six died, emphasize the severity of disease progression and the urgency to develop antivirals or an effective vaccine^1^. To that point, the World Health Organization discussed including EV71 and related D68 in its Blueprint List of Priority Diseases^1,2^.

Enterovirus 71 belongs to the *picornaviridae* family, and its genome consists of a positive-sense RNA approximately 7,500 nucleotides (nts) long that function to facilitate protein synthesis and virus replication^3,4^. EV71 regulates these events through several mechanisms, including recruitment of cellular proteins to specific regulatory sites located within the 5’ untranslated region (5’UTR). The 5’UTR is a dual-purpose RNA element that is predicted to fold into six stem loops (**Fig 1A**). Stem loop (SL) I facilitates genome replication, while stem loops II-VI promote cap-independent translation via a type I internal ribosome entry site (IRES)^4^. The IRES is essential to EV71 replication because it internally recruits the ribosome, which in turn drives synthesis of the complete enteroviral proteome.

**Fig.1.**
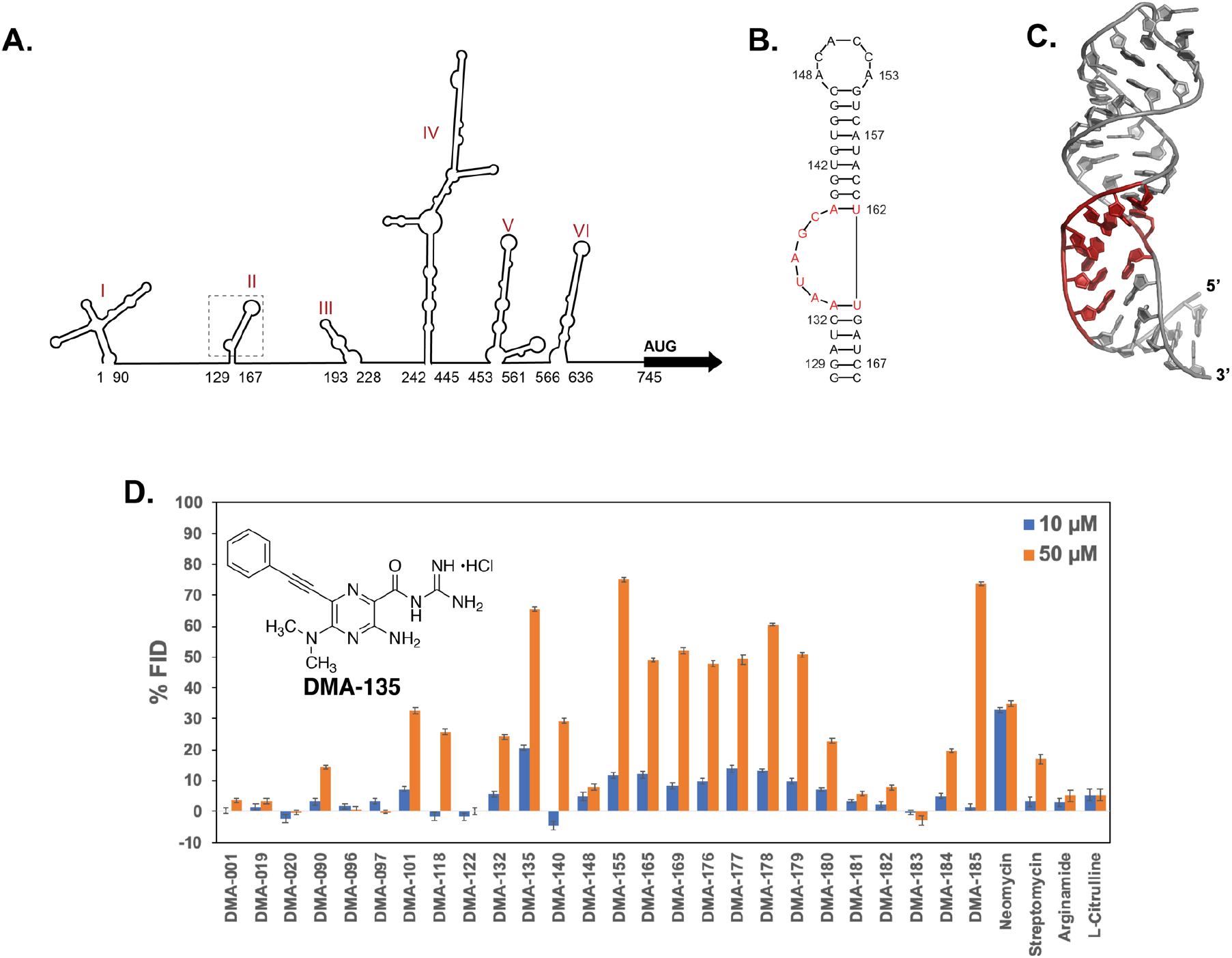
RNA-focused small molecules bind specifically to the EV71 SLII IRES domain. A) Schematic of the consensus secondary structure of the EV71 5′UTR determined using phylogenetic comparisons. (B) Secondary structure of EV71 SLII as validated by NMR spectroscopy^9^. (C) 3D structure (PDB code 5V16) of SLII refined against NOE derived distant restraints and RDCs^9^. (D) Screening data for small molecules vs the EV71 SLII (wt) IRES domain at 10 and 50 μM concentrations. Error bars shown are standard errors of mean calculated from three individual replicates. The chemical structure of DMA-135 is shown within the inset. Screening conditions were carried out in 50 mM Tris, 50 mM KCl, 0.01% Triton-X-100, 5% DMSO, pH = 7.4; Tat peptide: 50 nM; RNA: 90 nM; Small molecule: 10 or 50 μM; Excitation λ = 485 nm, Emission λ = 590 nm; Incubation time: 30 min; Error values shown are standard errors of mean calculated from three individual replicates.

Extensive studies of the EV71 IRES have revealed functional contributions of different RNA structures along with the collection of cellular proteins that interact with those structures^5–15^. Of particular importance, the stem loop II (SLII) domain interacts specifically with at least four cellular RNA binding proteins (RBPs) and a viral derived small RNA to modulate translation levels^5,7,9,10,16^. The secondary structure of the EV71 SLII domain folds into a phylogenetically conserved hairpin with a 7-nt apical loop and 5-nt bulge (**Fig 1B**)^9^. The three-dimensional structure of SLII shows that the apical loop and bulge form well-defined structural motifs (**Fig 1C**)^9^. Functional binding sites for the cellular hnRNP A1 and AUF1 proteins reside within the structured bulge, and these sites undergo a conformational change to form stable protein-RNA complexes^5,9^. HnRNP A1 binds the SLII bulge to stimulate translation whereas AUF1 competes for the bulge to counteract this stimulatory activity so that the levels of IRES-dependent translation are tuned to meet the replication needs of EV71. Genetic mutations or deletions of bulge residues inhibit EV71 replication, presumably by disrupting the protein-SLII regulatory axis that controls viral translation^9,16^. These collective observations provide strong support that the EV71 SLII domain, particularly the bulge region, is an attractive target to develop novel antiviral compounds.

Here, we screened an RNA focused small molecule library for binding to the EV71 SLII domain to identify five small molecule (SM) ligands that show specificity to the bulge when confirmed independently by NMR spectroscopy and titration calorimetry. One of the five SMs attenuates IRES-dependent translation and displays potent dose-dependent antiviral properties in cell culture. The NMR-derived 3D structure of the SM-SLII complex provides evidence that the small molecule binds to the bulge to induce a conformational change that fully exposes binding sites for AUF1 but not hnRNP A1. Calorimetric titrations of AUF1 into increasing molar ratios of the SM-SLII complex further validate that the small molecule increases the binding affinity of a ternary complex in a dose-dependent manner. This thermodynamic property of the SM-SLII-AUF1 complex was also independently observed both in an *in vitro* biochemical assay and in cell culture. When considered together, these results suggest that the SM acts through an allosteric mechanism to inhibit EV71 replication by stabilizing the translation repressive SM-SLII-AUF1 complex. Of note, the allosteric mechanism of action of the IRES targeting small molecule observed here for EV71 is fundamentally distinct from small molecules that target the HCV IRES^17–19^. In that case, benzimidazole inhibitors bind to the IRES subdomain IIa to induce a conformational change that leads to undocking of the IIb subdomain from the ribosome^19^. Thus, this study reports a novel mechanism of action of a small molecule that targets an IRES domain, and it provides a promising pathway to develop the first class of RNA-focused small molecule inhibitors of EV71 and likely the related EVD68.

## Results

### Identification of SM DMA-135 as an EV71 SLII ligand

To identify ligands for the EV71 SLII domain, a panel of RNA-targeted small molecules was screened against the native EV71 SLII domain using a fluorescent indicator displacement (FID) assay. As previously described this assay utilizes a fluorescently labeled, highly basic peptide fragment as an indicator that can be displaced from a wide range of RNA targets to assess small molecule binding^20^. We observed tight binding of the indicator peptide to SLII (24.5 ± 4.7 nM – **Figure S1**), which facilitated the development of a high-throughput FID protocol to screen small molecules against low concentrations of SLII (Z’ score 0.49, see SI). The small molecule library largely contained derivatives of amiloride, previously identified as a tunable RNA-binding scaffold^21,22^ (**Figure S2**), along with other known RNA binding small molecules. The screening assay was performed at two different small molecule concentrations to identify both strong and weak binding derivatives, and “hits” were defined as causing fluorescent changes greater than 25%. (**Fig 1C**). While 13 small molecule hits are observed at the higher (50 μM) concentration, only two prominent hits are observed at 10 μM: DMA-135, a C6 phenylacetylene substituted amiloride derivative, (**Fig 1C**) and a promiscuous RNA binder, neomycin. Quantitative binding analysis was then performed for five amilorides showing both significant displacement at 50 μM and structural diversity (**Fig 1C** and **S3**). This analysis confirmed that DMA-135 was the tightest SLII binder (CD_50_ = 12 μM, **Table S1**). While peptide assays for each RNA vary slightly, DMA-135 had been previously shown to have comparable affinity to HIV-1-TAR and HIV-2-TAR, much weaker binding to a model A-site and RRE-IIB RNAs^21^, and no interference from tRNA or DNA. We note that a direct measurement of *K*_d_ via ITC (see below) revealed a tighter binding constant (*K*_d_ ~ 500 nM) for EV71 SLII RNA. To further assess the nature of the highest affinity amiloride-SLII binders, NMR and calorimetric titrations were performed.

### Biophysical characterization of EV71 SLII-SM Interactions

We next used NMR titrations to locate the specific structural motifs on SLII that are recognized by the top five small molecule hits identified from the FID screen along with DMA-001, the core unsubstituted scaffold, as a control based on previous work^21,23^. For these experiments, we prepared an A(^13^C)-selectively labeled SLII construct as previously described^9^ and performed single point ^13^C-^1^H TROSY HSQC titrations. Figure 2A summarizes the effects that the addition of excess (5:1) small molecules have on the chemical shifts for each adenosine C8-H8 signal. Stem loop II contains eleven adenosines of which four are located within internal Watson-Crick base pairs (**Fig. 1B**). A133 and A139 are also involved in Watson-Crick base pairs; however, these residues are adjacent to the 5-nt bulge. None of the adenosines involved in internal base pairs (A130, A157, A159, and A165) or those located in the 7-nt apical loop (A148, A150, A153), show major chemical shift perturbations (CSPs) upon the addition of excess small molecules (**Fig. 2A**). By comparison, all six small molecules caused significant CSPs to adenosines adjacent to (A133 and A139) and located within (A134 and A136) the bulge. Thus, these data clearly localize the SLII binding sites for the small molecules to the surface formed by the bulge, in line with previous preferences for amiloride derivatives^21 22^(**Fig. 1B**).

**Fig.2.**
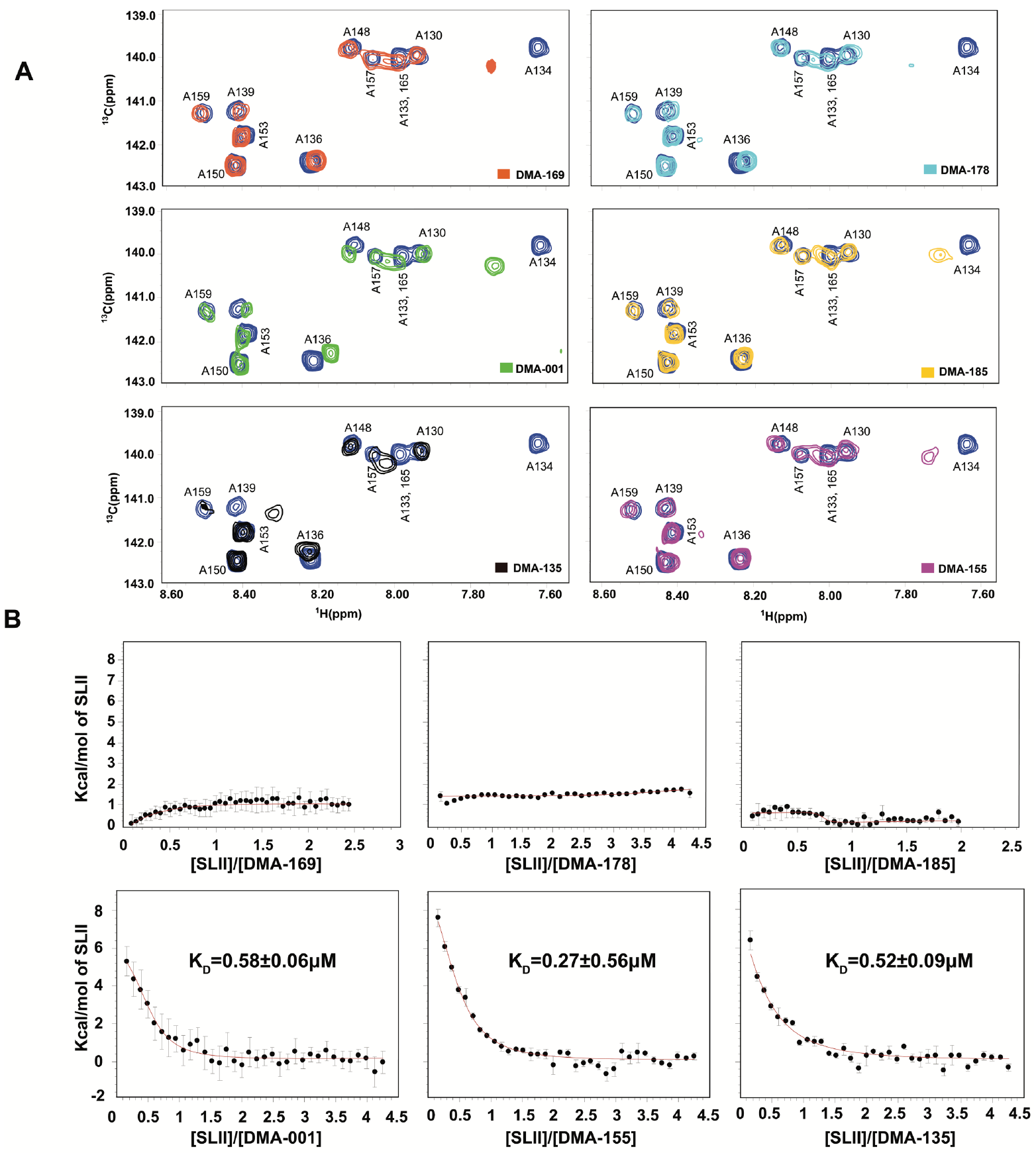
DMAs interact through the EV71 SLII bulge via an entropically driven mechanism. (A) Single-point ^1^H-^13^C TROSY HSQC titrations of A(^13^C)-selectively labeled SLII constructs with the respective DMAs added at 5-fold excess. The blue correlation peaks across each spectrum correspond to that of free SLII. The spectra were collected at 900 MHz in 10 mM K_2_HPO_4_, 20 mM KCl, 0.5 mM EDTA, and 4 mM BME (pH 6.5) D_2_O buffer at 298 K. (B) Calorimetric titrations of DMA-001, DMA-135 and DMA-155 reveal a single transition on the binding isotherm characterized by an entropically favored and enthalpically disfavored molecular recognition event. All titration data were processed and analyzed using Affinimeter^24^. The processed thermograms were fit to a 1:1 stoichiometric binding model, represented by the red lines. Reported values for dissociation constants (K_D_) and corresponding standard deviation are from triplicate experiments. Goodness of fit (*χ*^2^) of the experimental data to the 1:1 binding model are reported for each titration. The experiments were performed in 10 mM K_2_HPO_4_, 20 mM KCl, 0.5 mM EDTA, and 4 mM DTT (pH 6.5) buffer at 298 K.

To understand the thermodynamics of the SM-SLII interactions, we performed calorimetric titrations. The data reveal that only three (DMA-001, DMA-135 and DMA-155) of the six small molecules register a calorimetric response, and those three bind SLII with a favorable change in entropy and an unfavorable change in enthalpy (**Fig. 2B**). Fits to single-site binding models using Affinimeter^24^ showed excellent agreement with the experimental data therefore allowing reliable determination of the apparent binding constants, which ranged from ~300-600 nM (**Fig 2B**). Interestingly, the small molecule DMA-135 binds to SLII with moderately high affinity (K_D_ = 520 +/− 90 nM), yet it induces the most significant chemical shift perturbations observed in the HSQC titrations (**Fig. 2A**). These changes include the disappearance of the C8-H8 correlation peak belonging to A134, a large upfield shift of the H8 signal that belongs to A139, and other minor CSPs at residues A133, A136 and A165. By comparison, DMA-001 and DMA-155 induced smaller CSPs with the most notable difference at A134 where the addition of both SMs causes a downfield shift of the C8-H8 correlation peaks (**Fig. 2A**). In sum, the results indicate that all six small molecules interact with SLII through its bulge surface, the binding energies for three of the six SMs derive from large-favorable changes in entropy, and DMA-135 induces the largest changes to the SLII structure as determined by NMR titrations.

### DMA-135 attenuates EV71 translation and inhibits viral replication

Building on the specificity of the SM-SLII interactions, we next evaluated whether the small molecules affected EV71 IRES dependent translation in cells using a dual luciferase assay. The description of the bicistronic reporter plasmid, pRHF-EV71-5’UTR, used here has been described elsewhere^13^. In brief, the pRHF-EV71-5’UTR plasmid was used as a template to *in vitro* transcribe the dual luciferase reporter RNA (**Fig. 3A**), which was subsequently transfected into SF268 cells. Each SM was added to the cells at concentrations in the range of 0.001 – 1000 μM and after two days the activity of Renilla (RLuc) and Firefly (FLuc) were measured using the Promega dual-luciferase reporter assay. Only DMA-135 showed activity in the assay, and it attenuated IRES-dependent translation (FLuc) in a dose-dependent manner, without having any significant effects on cap-dependent (RLuc) translation (**Fig. 3B**). At 0.1 μM DMA-135, FLuc activity dropped by ~10% relative to the control. The FLuc activity was further reduced by 50% at 0.5 μM DMA-135 and no Firefly luciferase activity was detected at 100 μM DMA-135, even though Cap-dependent (RLuc) translation remained consistent at all concentrations (**Fig 3B**).

**Fig 3.**
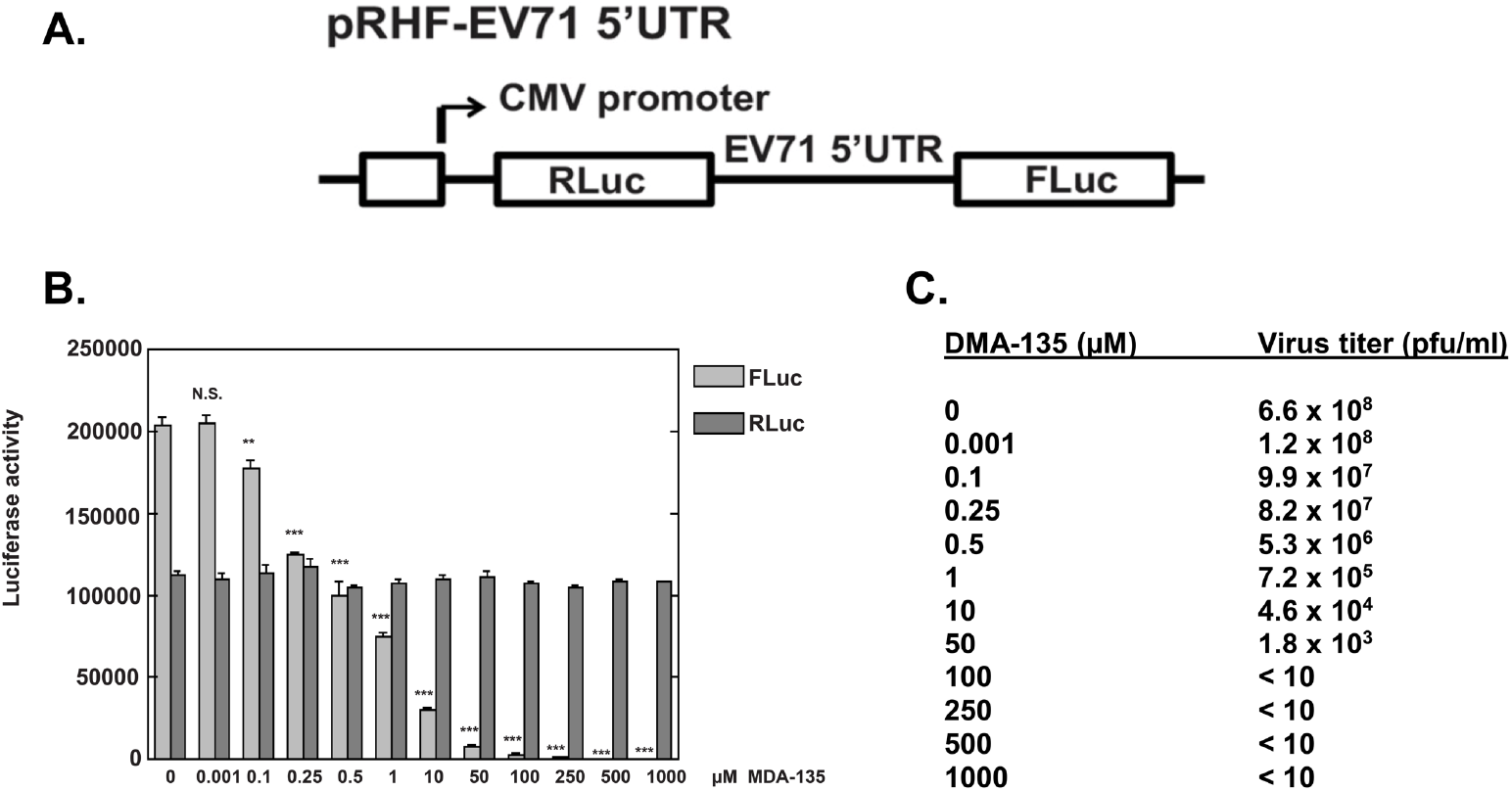
DMA-135 inhibits EV71 replication by abrogating IRES-dependent translation. A). The diagram depicts the bicistronic reporter plasmid used to synthesize RNA for transfections and dual-luciferase assays. (B) Effect of DMA-135 on EV71 IRES activity. SF268 cells were transfected with RLuc-EV71-5′UTR-FLuc RNA and cultured with various concentrations of DMA-135. Luciferase activity was measured 2 days later. Mean values and standard deviations from three independent experiments are shown in the bar graphs. ***P < 0.001; **P < 0.01; N.S., not significant. (C) Effect of DMA-135 on EV71 replication. SF268 cells were infected with EV71 at an moi = 1. Various concentrations of DMA-135 were added to the cells. Media were harvested 24 hr post-infection and assayed for infectious virus by plaque formation in Vero cells.

We next checked if DMA-135 had antiviral activity by infecting SF268 cells with EV71 at a multiplicity of infection (moi) of 1, followed by the addition of DMA-135. Harvested media were then applied to Vero cells and EV71 titers quantitated by plaque formation. Figure 3C shows that DMA-135 inhibits EV71 replication in a dose-dependent manner with a 2-log reduction in viral titers observed at 0.2 μM DMA-135 and a 5-log reduction when the concentration was increased to 50 μM. The IC_50_ of DMA-135 was determined to be 7.54+/−0.0024 μM and the CC_50_ in SF268 and Vero cells is >100 μM. Thus, the collective results indicate that DMA-135 inhibits EV71 replication by attenuating IRES-dependent translation at dosages of relatively low cellular toxicity.

### Solution structure of the SLII-(DMA-135) complex

To better understand the mechanism of action of DMA-135, we solved the NMR solution structure of the DMA-135-SLII complex. We previously reported the high-resolution 3D structure of free SLII (**Fig. 1C**), which was determined using a conjoined NMR-SAXS approach^9^. Informed by the collection of NMR data acquired there, we prepared a CU(^2^H),AG(^2^H_3’-5”_)-selectively labeled SLII construct to which DMA-135 was added at a 4:1 molar ratio. ^1^H-^1^H NOESY spectra collected on the SLII-(DMA-135) complex with this labeling scheme afforded detection of key intra-NOEs between A and G residues of SLII and inter-NOEs between the label sites on SLII and DMA-135. Figure 4 shows an overlay of the ^1^H-^1^H NOESY (*t*_m_=200 ms) spectra of free and the DMA-135 bound form of SLII. Comparisons of the NOESY spectra revealed interesting structural features of the SLII-(DMA-135) complex. First NOEs that involved the A133H2, A133H8 and A134H8 spin systems observed in free SLII were completely missing or highly attenuated in the complex (**Fig. 4A**). This implies that DMA-135 induces a local change in the structure of SLII that abrogates base stacking at positions 133-134. Second, the intra-NOE interactions that connected A136 to G137 and A139 to G140 were still present in the complex, albeit the chemical shifts of the corresponding H8 and H1’ signals were significantly different from those observed in free SLII. This indicates that the local structure at positions 136-137 and 139-140 are still preserved within the complex but that their chemical environments are different. Lastly, inter-NOEs between the methyl protons of DMA-135 and A133H8, A136H2, G137H8 and A139H2 firmly establish that DMA-135 forms site-specific contacts with the SLII bulge surface (**Fig. 4A**). As further validation of the DMA-135 complex, we synthesized a tri-fluorinated analog of DMA-135 (DMA-197) where a trifluoromethyl group was added to allow detection of the complex by ^19^F NMR. Figure 4B provides additional evidence of the (DMA-135)-SLII complex since the sharp 1D ^19^F NMR signal of DMA-197 shifts and becomes severely broaden when bound to SLII. We further verified that DMA-197 indeed binds the bulge surface on SLII by performing a single-point ^13^C-^1^H TROSY HSQC titration as described above for the other DMAs (**Fig. S4**).

**Fig.4.**
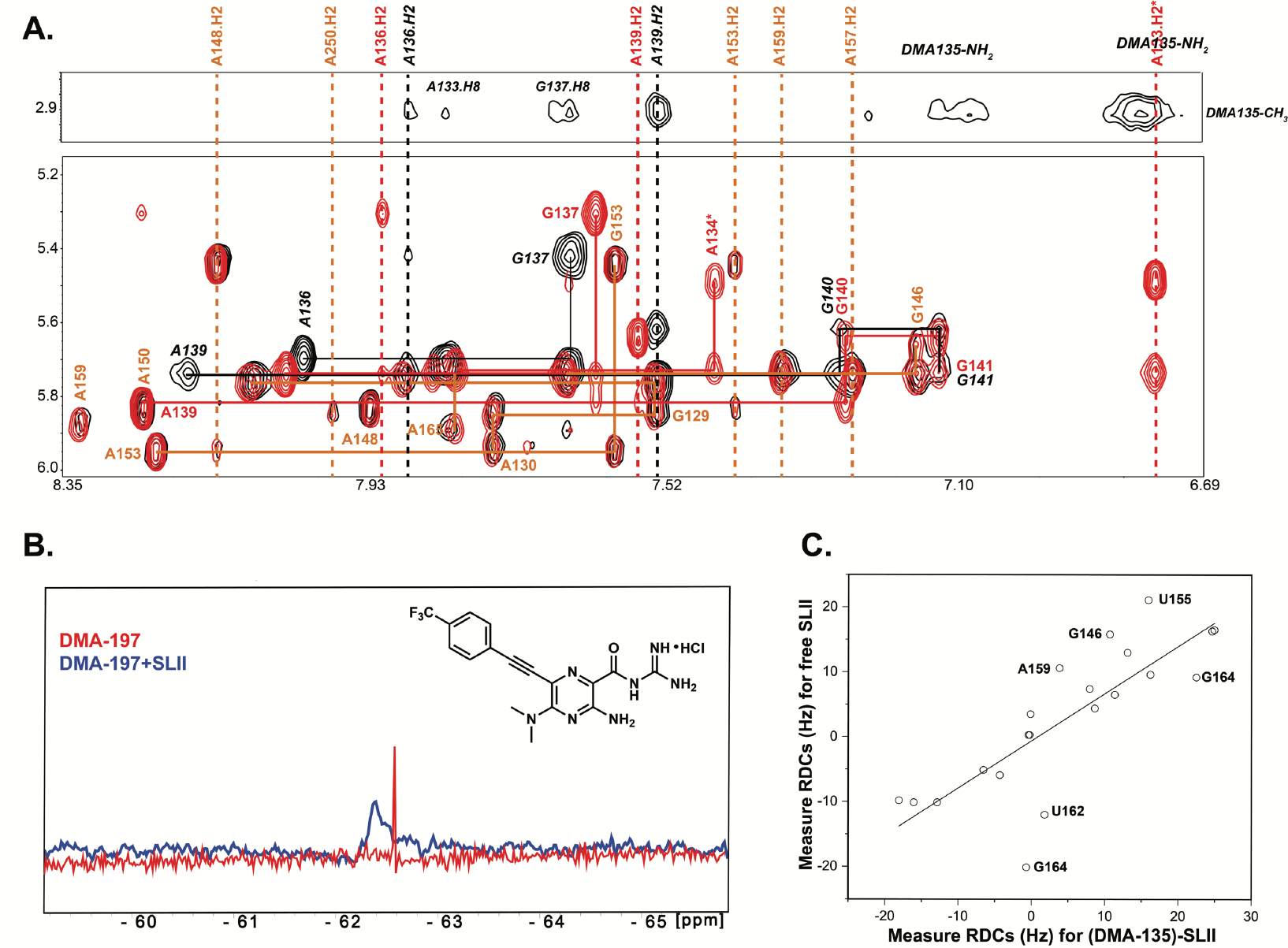
DMA-135 changes the local and global structure of the EV71 SLII IRES domain. (A) Overlay of ^1^H-^1^H NOESY spectra (t_m_=200 ms) of free SLII (red) and the (DMA-135)-SLII complex (black). The orange lines (solid and dashed) indicate NOE spin systems and chemical shifts that are identical between free SLII and the (DMA-135)-SLII complex. The black lines (solid and dashed) indicate NOE spin systems and chemical shifts of SLII within the complex that are significantly different from those of free SLII (indicated by red solid and dashed lines). Selected region of the ^1^H-^1^H NOESY spectrum of the (DMA-135)-SLII complex showing intermolecular and intramolecular NOEs between methyl protons on DMA-135 and nucleobases of SLII. (B) 1D ^19^F NMR spectra of DMA-197 in the absence (red) and presence of SLII at 1: complex (blue). (C) Correlation plot showing agreement between measured RDCs of free SLII and SLII within the DMA-135 complex. All NMR spectra were collected at 900 MHz in 10 mM K_2_HPO_4_, 20 mM KCl, 0.5 mM EDTA, and 4 mM BME (pH 6.5) D_2_O buffer at 298 K.

We proceeded to measure ^1^H-^13^C residual dipolar couplings (RDCs) of SLII in complex with DMA-135 so as to determine if DMA-135 also changes the global structure of SLII. Figure 4C shows that the correlation of measured RDCs between the free and bound forms of SLII are poor, indicating that DMA-135 also changes the global structure of SLII. We then calculated NMR structures of the SLII-(DMA-135) complex using NOEs and RDCs (see **Table S3**). Full details of the structure refinement routine are described in the Methods section. The ensemble of SLII-(DMA-135) structures reveal the same overall stereochemical features. First, DMA-135 binds through the bulge of SLII and induces a change in the local stacking patterns at positions 133-135 such that the bases are unstacked and dynamic within the context of the complex (**Fig. 5A** and **B**). Binding of DMA-135 to SLII also disrupts a Watson-Crick base pair formed between residues A133 and U163 (**Fig. 5B**). Second, A136 remains stacked on G137 as observed in free SLII and A139 also gives NOE patterns consistent with its remaining base paired to U162 (**Fig. 5B**). Because we were limited by inter-NOEs, the conformation of DMA-135 is dynamic within the ensemble so we cannot confidently describe specific interactions to SLII (**Fig. 5A**). However, we observed the appearance of new and broad NOEs within 6.7 – 7.1 ppm region of the SLII-(DMA-135) NOESY spectrum (**Fig. 4A**), consistent with the amino group of DMA-135 forming new hydrogen bonds within the context of the complex.

**Fig.5.**
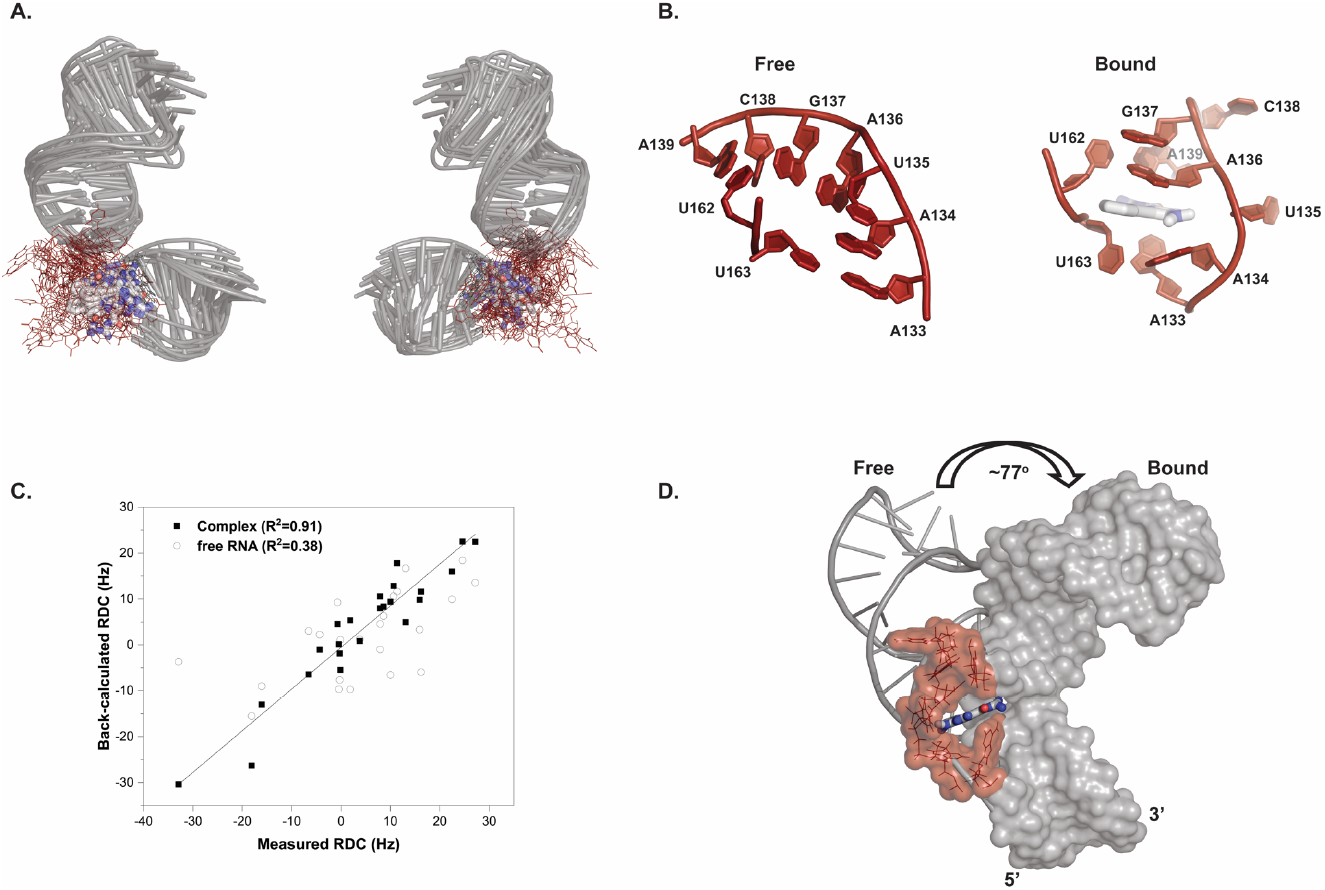
The structure of the EV71 SLII-(DMA-135) complex reveals that DMA-135 changes the local and global conformation of SLII. (A) Superimposition of 10 lowest energy SLII-(DMA-135) complex structures refined in AMBER using NOE-derived distance restraints and RDCs. The overall RMSD for the stably base paired regions is 2.4 Å. (B) Comparison of the bulge stacking interactions of (left) free SLII to that of the SLII-(DMA-135) complex (right). (C) Correlation plot between measured and back-calculated RDCs from a representative low energy SLII-(DMA-135) structure and to back-calculated RDCs from the free SLII structure. (D) Superimposition of the lower helices of free and bound SLII reveals that DMA-135 induces a global change to the SLII structure.

Since the SLII-(DMA-135) structure was refined with RDCs collected on the complex, we are able to observe that DMA-135 also induces a global change to the structure of SLII. This is evident by comparing the RDC refined structure of free SLII to that of SLII within the DMA-135 complex (**Fig. 5D** and **Supplementary video 1**). Figure 5D reveals that DMA-135 changes the inter-helical angle between the upper and lower helices of SLII by ~77°. This difference is consistent with the observation that DMA-135 changes the local stacking interactions of the bulge (**Fig. 5A**) such that this leads to a reorientation of the SLII helices within the complex. The interpretation of these collective observations is that DMA-135 binds to the bulge structure to induce a change in its stacking interactions, which exposes A133, A134 and U135 (**Fig. 5B**). By inducing the exposure of these residues, DMA-135 further changes the interhelical geometry of SLII (**Fig. 5D** and **Supplementary video 1**).

### DMA-135 allosterically increases the binding affinity of the AUF1-SLII complex in vitro and in vivo

The cellular proteins hnRNP A1 and AUF1 bind to the bulge of SLII to stimulate or repress IRES-dependent translation, respectively^5,9,13,16^. We therefore reasoned that DMA-135 might compete either or both of these interactions as part of its mechanism of action. To test this hypothesis, we performed calorimetric titrations using the tandem RNA binding domains (A1-RRM1,2 and AUF1-RRM1,2) of these proteins and the SLII-(DMA-135) complex prepared at a 5:1 molar ratio. Titrations of A1-RRM1,2 into the SLII-(DMA-135) complex showed that DMA-135 does not impact the binding reaction since the processed thermodynamic parameters are not significantly different when compared to that of the control titration (**Fig. S5A**). In contrast, excess DMA-135 increased the apparent binding affinity of AUF1-RRM1,2 for SLII by ~3-fold (**Fig 6A**). We further showed that this property is dose-dependent since the binding dissociation constant decreases as the molar ratio of (DMA-135):SLII increases (**Fig 6B**). This property is also specific to DMA-135 because we did not observe significant differences in binding affinities of AUF1-RRM1,2 for SLII when SLII is pre-equilibrated with excess (5-fold) DMA-001 or DMA-155 (**Figs. 6A** and **S4B**). Together, these experiments clearly demonstrate that the effects of DMA-135 on SLII-protein binding are specific to AUF1-RRM1,2 in a dose-dependent manner.

**Fig.6.**
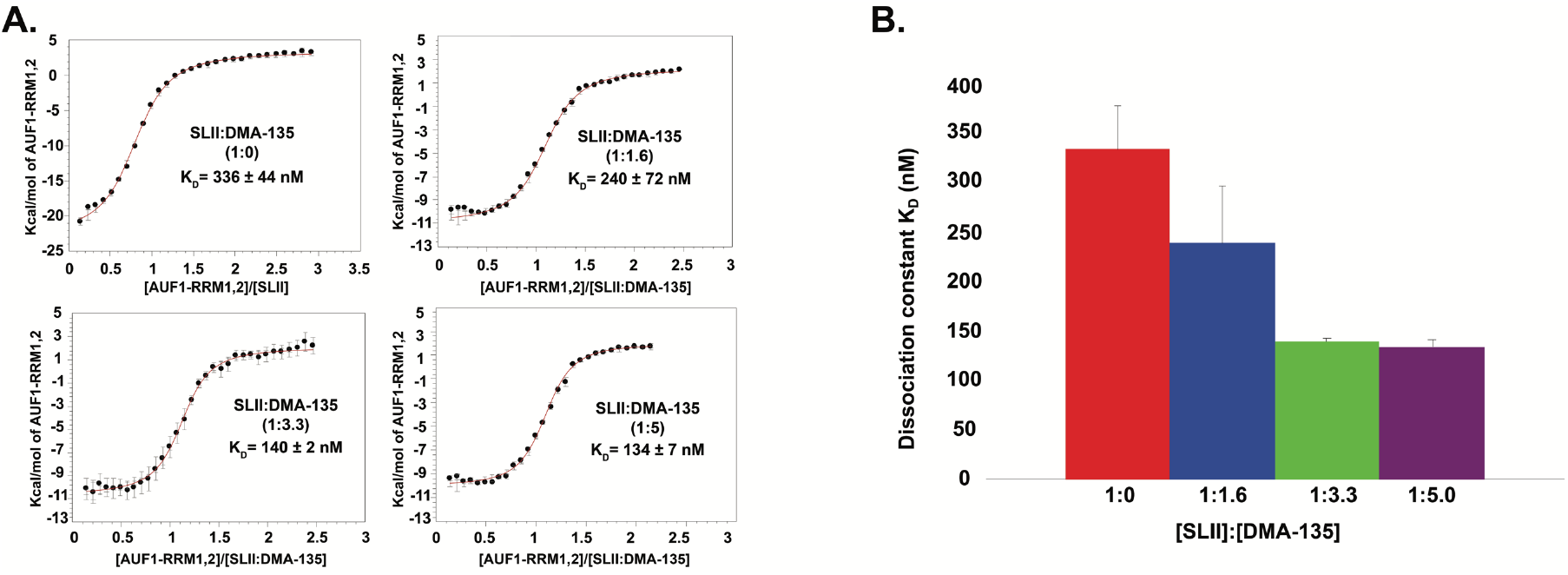
DMA-135 allosterically increases the affinity of AUF1 for the EV71 SLII IRES domain via the formation of a ternary complex. (A) Calorimetric titrations of AUF1-RRM1,2 to SLII or to SLII:DMA-135 complex at different molar ratios reveal a single transition in the binding isotherm characterized by and enthalpically driven molecular recognition event. All data were processed and analyzed using Affinimeter^24^. The processed thermograms were fitted to a 1:1 stoichiometric binding model, represented by the red lines. Values for dissociation constants (K_D_) and corresponding standard deviation are reported for each titration. The experiments were performed in 10 mM K_2_HPO_4_, 20 mM KCl, 0.5 mM EDTA, and 4 mM BME (pH 6.5) buffer at 298 K. (B) Graphical representation of the dissociation constants for the AUF1-RRM1,2-SLII complex as a function of DMA-135 concentration. Reported values for K_D_ values and corresponding standard deviations are from triplicate experiments.

Using an ^15^N-labeled AUF1-RRM1,2 construct, we verified by NMR spectroscopy that DMA-135 does not interact with isolated AUF1-RRM1,2 since the ^15^N-^1^H HSQC peaks are identical to those collected without excess DMA-135 (**Fig S6A**). Instead, we observed the appearance of a new set of correlation peaks when the SLII-(DMA-135) complex is titrated into ^15^N-labeled AUF1-RRM1,2 (**Fig S6A**). These correlation peaks are not observed in the spectrum recorded when only SLII is titrated into AUF1-RRM1,2 (**Fig S6A**). When considered together, the calorimetric titrations and NMR data provide evidence for the formation of a (AUF1-RRM1,2)-SLII-(DMA-135) ternary complex (**Fig S6B**).

To test whether DMA-135 affects the stability of the AUF1-SLII complex in a biological context, we designed a set of *in vitro* and *in vivo* biochemical experiments. In the *in vitro* assay, biotinylated SLII was exposed to SF268 cell lysate in the presence of various concentrations of DMA-135. Complexes of AUF1-SLII were pulled down by streptavidin and the abundance of AUF1 was determined by Western blotting. Figure7A reveals that the levels of AUF1 increase in the pulldowns as the concentration of DMA-135 increases, indicating that the addition of DMA-135 promotes binding of AUF1 to SLII as observed by calorimetry (**Fig 6A**). To investigate the effects of DMA-135 in a cellular context, biotinylated wild type SLII and a CCC mutant that abrogates AUF1 and hnRNP A1 binding^5,16^ were transfected into SF268 cells. Cell lysates were prepared 24 hr after transfection and the levels of AUF1 being pull down were determined by Western blotting. Figure 7B shows that the addition of DMA-135 increases binding of AUF1 to wild-type SLII but not to the CCC mutant, consistent with the interpretation that DMA-135 promotes the formation of a specific ternary complex with the wild-type RNA.

**Fig 7.**
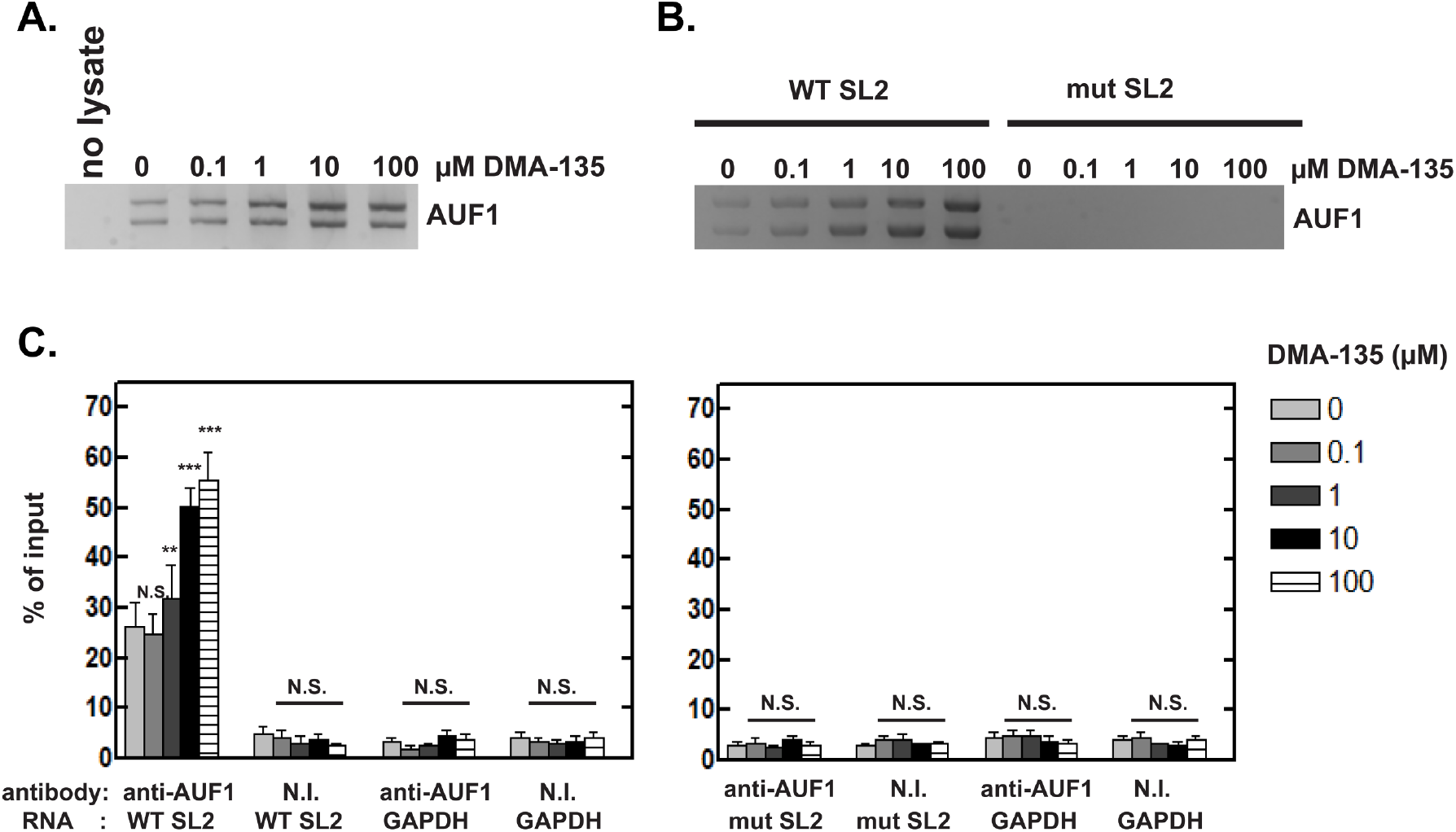
Allosteric effects of DMA-135 observed within the context of the cellular environment. (A) Protein-biotinylated RNA pull-down experiments were performed to examine the effect of DMA-135 on the interaction between AUF1 and the EV71 SLII IRES domain. Biotinylated SLII was incubated with SF268 cell lysate to which was added various concentrations of DMA-135. Streptavidin-linked beads were used to pull-down biotin-labeled SLII and its associated cellular proteins. Samples were analyzed by Western blot using anti-AUF1 antibody. (B) Biotinylated wild-type SLII or the CCC-bulge mutant SLII were transfected into SF268 cells. Cells were cultured with various concentrations of DMA-135. Cell lysates were used for pull-down assays for SLII-associated proteins as described above. AUF1 bound to SLII RNA was detected by Western blotting. (C) RIP assays for AUF1-EV71 Luc RNA interactions in cells. SF268 cells were transfected with RLuc-EV71 5’UTR-FLuc RNA, harboring the wild-type SLII, or with CCC-bulge mutant SLII and cultured with various concentrations of DMA-135. Lysates were prepared and analyzed by ribonucleoprotein immunoprecipitation (RIP) with non-immune antibody or AUF1 antibody. Both input and immunoprecipitated materials were analyzed by qRT-PCR for EV71 RNA. NI, non-immune antibody. The means ± standard deviation (SD) from three independent experiments are shown. ***P < 0.001; **P < 0.01; N.S., not significant.

Lastly, we performed RNA-protein immunoprecipitations followed by qRT-PCR to assess whether DMA-135 affects the association of AUF1 with the EV71 5’UTR in cells. SF268 cells were transfected with bicistronic reporter RNAs (**Fig 3A**) containing wild type SLII or the CCC mutant in the presence of increasing concentrations of DMA-135. Cell lysates were prepared 24 hr post transfection and RNA-protein complexes were immunoprecipitated using non-immune antibody or AUF1 specific antibody. RNA isolated from immunoprecipitates was then analyzed by qRT-PCR. Wild-type EV71 5’UTR was enriched by antibody against AUF1 relative to non-immune antibody, and the addition of DMA-135 increased the levels of wild-type EV71 5’UTR pulled down by AUF1. The EV71 5’UTR containing the CCC mutation in SLII was not detected by qRT-PCR, consistent with the observation that this mutation abrogates AUF1 binding. The Western blot in Figure S6 demonstrates that the AUF1 antibody precipitated AUF1 while the non-immune antibody did not. Thus, the independently designed *in vitro* and *in vivo* biochemical assays validate that DMA-135 functions to allosterically enhance the binding of AUF1 to SLII by forming a specific ternary complex.

## Discussion

Given the abundance of functional RNA structural elements in human disease systems and in pathogens that infect humans, RNA structures represent significant untapped targets in drug discovery. Here, we report a comprehensive study that combined small molecule screening, biophysics and virological assays to successfully identify a small molecule (DMA-135) that targets the IRES domain of EV71 to inhibit its replication in cell culture. Characterization of DMA-135 activity revealed that it acts as an allosteric inhibitor to increase the binding affinity of AUF1 for the SLII IRES domain and suggested that a ternary complex consisting of AUF1-SLII-(DMA-135) (**Fig S3B**) attenuates IRES-dependent translation. Our results thus support the targeting of RNA structures of relatively moderate complexity with small molecules and that the mechanisms of inhibition include allosteric effects.

In this study, we chose to target the type I IRES domain of enterovirus 71, which, along with the related D68, poses serious threats to human health, particularly in regions of Southeast Asia, where it also challenges the economy^1,25–28^. To date, there is no broad-spectrum vaccine or antivirals to slow EV71 progression. The type I IRES domain represents an attractive target for therapeutic intervention since it folds into a highly structured RNA that interacts with multiple cellular proteins to regulate protein synthesis; however, detailed structure-function knowledge is lacking for most of these interactions. Stem loop II is an exception as its sequence conservation and its different binding partners have been thoroughly characterized^5–7,10,13,16^. Those studies demonstrated that both the sequence and structure of the SLII bulge surface are essential for EV71 replication, because genetic mutations or deletions to its residues abrogate a network of interactions that modulate viral translation^9,16^. With that knowledge, we reasoned that small molecules might also disrupt the SLII regulatory axis.

The structure of the SLII-(DMA-135) complex and the thermodynamic description of its formation revealed that DMA-135 binds to the shape created by the bulge surface and induces a change that makes a subset of the residues more dynamic (**Fig. 5**). The residues that transition from a stacked to an unstacked conformation include A133, A134, and U135. These residues match part of the high affinity consensus sequence motif (AU-rich elements) recognized by AUF1^29–31^. The combination of structural changes and increased binding affinity of AUF1 to SLII indicate an allosteric mechanism of action of DMA-135. Support for this mechanism was also observed within the context of the cellular environment. Particularly, a dose-dependent increase in binding of AUF1 to the EV71 5’-UTR was observed at DMA-135 treatment concentrations that strongly inhibit viral translation and viral replication.

An additional feature of DMA-135 was that it changed the global architecture of SLII such that the lower and upper helices shift their relative orientations within the complex (**Fig 5D**). Since we do not know how the SLII structure fits within the larger IRES or how the IRES coordinates ribosome assembly, we cannot confidently assign a function to the relative helical orientation adopted due to base stacking within the bulge. Nevertheless, we previously showed that mutating the central 5’-UAG-3’ bulge sequence to 5’-CCC-3’ also changes the relative helical orientation and this mutant inhibits EV71 replication by attenuating translation^9^. Therefore, this might be a secondary aspect of the mechanism of action of DMA-135.

While there are reports demonstrating binding of small molecules to the HCV IRES that result in structural changes^17,19^, it is important to note that different IRES types possess different structures and mechanisms of action. The HCV IRES is type III, and the EV71 IRES is type I. The HCV IRES promotes internal initiation of translation by direct interaction with the 40S ribosomal subunit. The EV71 IRES acts via binding eukaryotic initiation factors and host RNA-binding proteins. The IRES structures of HCV and EV71 are also structurally distinct. The EV71 IRES is composed of six stem-loops in which numbers II-VI comprise the IRES. By contrast, the HCV IRES is composed of four stem-loops with no structural homology with EV71. HCV does not possess a structure similar to EV71 SLII^32,33^, and thus the mechanism of action of the (DMA-135)-SLII complex is novel relative to small molecules that target the HCV IRES. Moreover, the small molecules that target HCV appear to function by changing how the IRES binds the 40S subunit^19^. Our work shows a novel mechanism of action by which DMA-135 functions to allosterically recruit AUF1, which in turn down-regulates EV71 translation.

In sum, we report a thorough study of a small molecule-RNA complex that inhibits EV71 replication by attenuating translation. Through the process, we also discovered that the small molecule acts as an allosteric inhibitor through the formation of a ternary SM-RNA-protein complex. We also note that, while amiloride derivatives have been shown to interact modestly with other RNAs such as HIV-1-TAR, DMA-135 has not been shown to have biological activity prior to this work, and that the combination of unique small molecule-induced RNA structural changes that lead to the allosteric recruitment of AUF1; specific biological activity of DMA-135; and its lack of toxicity in these studies reflect a high level of functional selectivity. This assessment is further supported by the close agreement between the measured binding affinities and biological IC_50_ values. We believe that this study serves as strong evidence to support drug discovery efforts intended to target RNA structures while revealing new approaches and potential mechanisms of action. Along those lines, ongoing efforts include optimizing the binding affinity of DMA-135 and also evaluating its utility as a broad-spectrum inhibitor against the related EVD68, which also contains a phylogenetically conserved SLII domain.

## Methods

### RNA preparation

EV71 stem loop II was prepared by *in vitro* transcription using recombinant T7 RNA polymerase that was overexpressed and purified from BL21 (DE3) cells. Synthetic DNA templates, corresponding to the EV71 2231 isolate, were purchased from Integrated DNA Technologies (Coralville, IA). Transcription reactions were performed using standard procedures and consisted of 3-6 mL reaction volumes containing unlabeled rNTPs, (C^13^/N^15^)-labeled rNTPs or ^2^H-labeled rNTPs wherein positions 3’-5’’ and H5 are deuterated. Following synthesis, samples were purified to homogeneity by 10% denaturing PAGE, excised from the gel, electroeluted, and desalted via exhaustive washing of the samples with RNase-free water using a Millipore Amicon Ultra-4 centrifugal filter device. Samples were annealed by heating at 95 °C for 2 minutes and flash-cooled on ice. Samples were subsequently concentrated to desired NMR levels using a Millipore Amicon Ultra-4 centrifugal filter device, followed by addition of buffer salts only [5 mM K_2_HPO_4_ (pH 6.5)] or with buffer salts plus the addition of 0.5 mM EDTA, 20 mM KCl, and 4 mM BME. Concentration of samples were determined using the RNA theoretical molar extinction coefficient, and NMR samples ranged from 0.05 to 0.2 mM at 200 μL. RNA for all titration experiments was annealed as described above. Post-annealing, all RNAs were washed into their respective buffers (see below), using a Millipore Amicon Ultra-4 centrifugal filter device.

### Small molecule synthesis and library preparation

Synthesis of amiloride derivatives DMA-001, DMA-019, DMA-096, DMA-097, DMA-101, DMA-132, DMA-135, DMA-155, DMA-165, DMA-169, DMA-178, and DMA-185 have been previously reported^21,22^. 4-Trifluoromethyl derivative DMA-197 was synthesized via the previously reported 2 step procedure using Sonogashira coupling reaction followed by guanidinylation^21^. Structures of small molecules studied along with the characterization data for the new derivative DMA-197 presented in the supplementary information. Other small molecules studied – neomycin, streptomycin, mitoxantrone, Hoechst 33258, argininamide, and L-citrulline were purchased from commercial suppliers and used as is. All small molecules except neomycin and streptomycin were dissolved in DMSO at a stock concentration of 50 mM and used in assays as described below. Neomycin and streptomycin are insoluble in DMSO and hence dissolved in water at 50 mM concentration.

### Binding assay for indicator peptide to EV71 SLII RNA

Binding constant of the fluorescently labeled peptide for EV71 SLII RNA, was measured using previously reported procedure^20^. Briefly, a Tat-derived peptide [N-(5-FAM)-AAARKKRRQRRRAAAK(TAMRA)-C] was purchased from Lifetein and dissolved at a concentration of 120 nM in the assay buffer containing 50 mM Tris, 50 mM KCl, 0.01% Triton-X-100, pH = 7.4. RNA stock solutions were serially diluted in assay buffer from 0 - 2 μM concentrations. The Tat peptide solution (8 μL) and RNA solution (8 μL) were combined in a Corning™ low volume round-bottom black 384 well-plates, shaken on an orbital shaker for 5 min, centrifuged at 4000 rpm for 1 min, and allowed to incubate for 30 min (Final Tat concentration = 60 nM, final RNA concentrations = 0 - 1000 nM). The 384 well-plates were then read on a Clariostar™ monochromator (BMG Labtech) microplate reader using the excitation and emission wavelengths of 485 nm (FAM) and 590 nm (TAMRA) respectively. The fluorescence intensities at each RNA concentration were normalized to the Tat-only control wells. Dissociation constants (*K*_d_) were calculated by fitting the observed relative fluorescence intensity values at each concentration to the equation below using the GraphPad Prism curve fitting software. Assays were conducted in triplicates and each replicate contained three technical replicates.

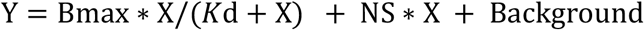

Where, Y = Total binding; X = Conc. of added ligand; Bmax = Maximum binding; *K*_d_ = Equilibrium dissociation constant; NS: Slope of Nonlinear regression; Background: Measured binding without the ligand.

### Small molecule screening using the Tat peptide displacement assay

Screening of the small molecule library against the EV71 SLII wt RNA was performed using the Tat peptide displacement assay previously reported^20^. Briefly, Tat peptide was diluted at a concentration of 180 nM in the assay buffer. RNA solution was prepared by diluting the stock solution in the assay buffer at 270 nM concentration. These concentrations were chosen to provide the best fluorescence intensity possible in the bound state. Small molecules were diluted to concentrations of 30 μM and 150 μM. Tat peptide (6 μL), RNA (6 μL), and small molecule solution (6 μL) were combined in a 384 well-plate. Final concentrations in each assay well were as follows-Tat peptide: 60 nM; RNA conc: 90 nM; small molecule: 0 μM, 10 μM, and 50 μM. The well-plate was shaken on an orbital shaker for 5 min, centrifuged at 4000 rpm for 1 min, and allowed to incubate in dark for 30 min and read using the excitation and emission wavelengths of 485 nm (FAM) and 590 nm (TAMRA) respectively. % Displacement of Tat peptide from RNA was calculated using the equation shown below. Each screening experiment was performed in triplicates and each replicate contains a technical triplicate.

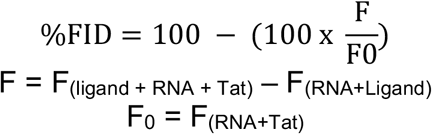

Molecules that show >20% FID at tested concentrations of 10 μM or 50 μM, were labeled as hits of the screening assay at that concentration.

### Full binding titrations of amiloride derivatives with EV71 SLII

Titrations of amiloride derivatives for binding to both the wild type and mutant RNAs was performed as previously described^3,20^. Briefly, small molecules were diluted in the assay buffer to achieve final assay concentrations between 0-334 μM. Tat peptide (6 μL), RNA (6 μL), and small molecule solution (6 μL) were combined in a 384 well-plate, shaken on an orbital shaker for 5 min, centrifuged at 4000 rpm for 1 min, and allowed to incubate for 30 min before being read using the excitation and emission wavelengths of 485 nm (FAM) and 590 nm (TAMRA) respectively. Observed fluorescence at each small molecule concentration was normalized to the fluorescence of Tat peptide: RNA complex. Competitive displacement of 50% of peptide from RNA was calculated by fitting the normalized data to the equations S2 and S3 below using the GraphPad Prism curve fitting software. Assays were conducted in triplicate and each replicate contained three technical replicates. Errors presented are the standard errors of mean from three replicates.

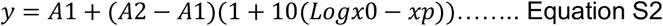

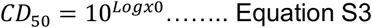

### Protein Purification

The AUF1 protein used in this study (residues from 70 to 239) was subcloned into the pMCSG7 vector^34^; subsequently overexpressed as an N-terminal (His)_6_-tagged fusion protein in BL21(DE3) cells and grown in either LB Broth or M9 minimal medium supplemented with ^15^NH_4_CL (0.5 g/L). The (His)_6_-tagged AUF1 construct was purified via nickel affinity chromatography on a Hi-trap column (GE Biosciences) followed by a Hi-trap Q column (GE Biosciences). The (His)_6_-purification tag was cleaved using the TEV enzyme and the cleavage mixture was then loaded onto Hi-trap columns (GE Biosciences) to isolate the protein of interest. Subsequently, the AUF1 construct was loaded onto a HiLoad 16/600 Superdex 75 pg (GE Bioscience) gel filtration column and eluted into the desired buffer. Protein stock solutions were kept in a buffer consisting of 10mM K2HPO_4_, 0.5 mM EDTA, 20 mM KCl, and 4 mM BME (pH6.5). Protein homogeneity and purity was confirmed by SDS-PAGE, and concentration was determined using the theoretical molar extinction coefficient. C-terminal (His)_6_-tagged UP1 (A1-RRM1,2) was prepared as previously described^9^.

### Isothermal Titration Calorimetry Experiments

The RNA and the respective DMA samples were prepared and purified as described above. Calorimetric titrations were performed on a VP-ITC calorimeter (Microcal, LLC) at 25 °C into 10 mM K_2_HPO_4_, 20 mM KCl, 0.5 mM EDTA, and 4 mM BME (pH 6.5) buffer, centrifuged and degassed under vacuum before use. SLII at 40 μM was titrated into ~1.4 mL of 1.5 μM of the respective DMA over a series of 32 injections set at 6 μL each. To minimize the accumulation of experimental error associated with batch-to-batch variation, titrations were performed in duplicate. Data were analyzed using KinITC routines supplied with Affinimeter^24^.

The AUF1 construct used in this study was prepared and purified as described above. Calorimetric titrations were performed on a VP-ITC calorimeter (Microcal, LLC) at 25 °C into 10 mM K_2_HPO_4_, 20 mM KCl, 0.5 mM EDTA, and 4 mM BME (pH 6.5) buffer, centrifuged and degassed under vacuum before use. AUF1 at 100 μM was titrated into ~1.4 mL of 8 μM SLII:DMA-135 complex at the following RNA:DMA ratios: 1:0, 1:1.6, 1:3.3, 1:5. To minimize the accumulation of experimental error associated with batch-to-batch variation, titrations were performed in duplicate. Data were analyzed using KinITC routines supplied with Affinimeter.

### Cells and virus

SF268 (human glioblastoma) and Vero (African green monkey kidney) cells were cultured as described^16^. Cells were infected with EV71 (TW/2231/98) at indicated moi (multiplicity of infection) and incubated 1 hr at 37°C for adsorption. Unbound virus was removed, and cells were refed fresh medium with various concentrations of DMA-135. Media from infected cells were harvested 24 hrs post-infection, and virus titers were determined by plaque assay on Vero cells.

### Determination of EV71 IRES activity

The bicistronic reporter plasmid pRHF-EV71-5′UTR which contains the EV71 IRES between Renilla and Firefly luciferase open reading frames and the synthesis of RLuc-EV71-5′UTR-FLuc RNA were described previously^13^. SF268 cells were seeded in 24-well plate. Two hundred ng of reporter RNA, 5μl of SuperFect (Qiagen), and 400μl of RPMI with 10% FCS were combined and added to one well of cells. Cells were incubated at 37°C for 4hrs and medium were changed and various concentrations of DMA-135 were added. Two days after transfection IRES activity was determined by measuring Renilla luciferase (RLuc) and Firefly luciferase (FLuc) activities using a dual-luciferase reporter assay system (Promega).

### Plaque reduction assay

Anti-viral activity of DMA-135 was determined by plaque reduction assay. In brief, Vero cells were infected with EV71 at a concentration of 10^7^ pfu/ml which gives approximately 60 plaques per well. DMA-135 was diluted and incubated in the agar-medium overlay at 37°C for 3 days. Cells were stained and plaques were count. The concentration of DMA-135 required to reduce the number of plaques by 50% relative to the virus control was expressed as IC_50_. Experiments were performed in triplicate.

### Cytotoxicity assays

Various concentrations of DMA-135 were added to SF268 and Vero cell lines. The cells were incubated at 37°C for 96 hrs. Cell viability was determined by MTT assay and measured at 570 nm according to the manufacturer’s instructions (EMD Millipore). All experiments were performed in triplicate. The concentration of DMA-135 required to reduce cell viability to 50% of the control cells was expressed as CC_50_.

### Pull-down of protein-biotinylated RNA complexes using streptavidin beads

SF268 cell extracts were prepared as described^5^. Biotin-labeled CMV-RLuc-EV715’UTR (wild type or CCC mutant)-FLUC and biotin-labeled SL2 (wild type or CCC mutant) were synthesized using biotin-16-UTP (Roche). Binding reaction and pull-down were carried out as described^5^. The eluted proteins were fractionated by 10% SDS-PAGE. AUF1 was detected by Western blot analyses using anti-AUF1 rabbit polyclonal antibody at 1: 15,000 dilution.

### RNP-immunoprecipitation

Immunoprecipitations of endogenous protein-RNA complexes were used to examine association of AUF1 with EV71 5’UTR in SF268 cells as described^5^. For quantitations of mRNAs in precipitates, purified RNAs were reverse transcribed into cDNAs using the High Capacity cDNA Reverse Transcription kit (Applied Biosystems), followed by SYBR green quantitative PCRs. The forward and reverse primer sequences, respectively, are as follows: EV71: 5’-CCCACCCACAGGGCCCACTGG-3’ and 5’-CGTTGATTTACAGCTTCTAA-3’; GAPDH: 5′-TTTAACTCTGGTAAAGTGGATATTGTTG-3′ and 5′-ATTTCCATTGATGACAA GCTTCC-3′.

### NMR data acquisition

NMR spectra were recorded on Bruker Advance (700, 800 and 900 MHz) high-field spectrometers. ^1^H-^13^C HSQC titrations were collected in 100% D_2_O at 303 K on SLII constructs with selectively labeled ^13^C(rATP). ^1^H-^1^H NOESY (t_m_=200 ms) spectra of the (DMA-135)-SLII complex were collected in 100% D_2_O at 303 K on SLII samples prepared with selectively deuterated rNTPs, rRTP ^2^H (3’,4’,5’5’’) and rYTP^2^H (5,3’,4’,5’5’’). Residual dipolar couplings (RDCs) were measured using ^1^H-^13^C TROSY-HSQC experiments for both isotropic and anisotropic ^13^C-selectively labeled SLII samples. Partial alignment was achieved via the addition of Pf1 filamentous bacteriophage (ASLA) to a concentration of ~15mg/mL; phage concentration was validated via ^2^H splitting at 900 MHz. RDC values were determined by taking the difference in ^1^J_CH_ coupling under anisotropic and isotropic conditions. ^1^H-^15^N HSQC titrations of either unlabeled SLII, DMA-135 or SLII:DMA-135 complex into ^15^N-labeled AUF1 were performed. Spectra were collected at the following molar ratios SLI^I^:AUF1 (1:1), DMA-135: AUF1 (4:1), SLII:DMA135: AUF (1:4:1). All NMR data were processed with NMRPipie/NMRDraw^35^ and analyzed using NMRView J^36^ or Sparky^37^.

The amiloride DMA-135 protons were assigned using 1D proton, 1D carbon and NOESY NMR experiments. Spectra were collected at a concentration of 200 *μ*M of the small molecule dissolved in D2O. The dimethylamino group protons were observed at a chemical shift of 3.00 ppm.

### HADDOCK-derived starting models of the (DMA-135)-SLII complex

Since the ^1^H-^1^H NOESY data (**Fig. 4**) revealed that DMA-135 abrogates base stacking interactions within the SLII bulge, we first performed molecular dynamic simulations on the NMR refined SLII structure (PDB code = 5V16) using distance restraints with lower bounds set at 7 Å for bulge residues A133-U135. This simulation resulted in an ensemble of SLII structures where the bulge loop was remodeled so as to disrupt local base stacking interactions at residues A133-U135. We next performed HADDOCK^38^ (version 2.2 Guru mode) docking calculations with these remodeled structures and incorporated the chemical shift perturbations as well as the inter-NOEs observed between SLII and DMA-135 as active restraints. The bulge loop (residues A133-C138) was treated as semi-flexible. One thousand rigid body (DMA-135)-SLII complex structures were calculated, and the most favorable structures (n=195) underwent further semi-flexible and water refinement. The output structures were clustered with a minimum size of 4, resulting in 7 total clusters. Sixteen representative (DMA-135)-SLII complex structures from the three best-scoring HADDOCK clusters were selected for further refinement in AMBER^39,40^. The Molecular Operating Environment (MOE) was used to generate the initial structure for DMA-135. Protonated DMA-135 was processed through PRODRG2 server.

### AMBER equilibration and simulation workflow

Sixteen starting (DMA-135)-SLII complex structures were prepared for AMBER using Antechamber and tLEaP. Initial equilibration of these systems was performed through a series of energy minimizations. Final frames of the minimization steps were simulated using pmemd.cuda for 50 ns each in AMBER 16 using the ff99bsc0_OL3_ force field. In both minimization and production, simulations were performed in implicit solvent using generalized Born model (igb = 1) and a salt concentration of 10 mM.

### Details of the Antechamber step

Antechamber and QM calculations were utilized to build input files optimized for simulation in AMBER. The parameters for the DMA-135 were obtained using the GAFF force field and the Antechamber package. DMA-135 structures were prepared for Antechamber and further AMBER calculations through sqm (QM) calculation of atomic point charges and the reduce program. Antechamber was performed on each structural model, assuming a net charge of +1 due to a protonated guanidinium group. An additional force field containing specific parameters was written for each small molecule (SM)-RNA construct using the parmchk feature of Antechamber. The output files of Antechamber calculations were utilized to perform two rounds of tLEaP. First, GAFF and ff14SB were used to generate AMBER parameter files for DMA-135, and a library (*.lib file) was created. Second, the ff99bsc_OL3_ and ff14SB were sourced and the aforementioned library loaded, allowing the generation of AMBER parameters (*.prmtop and *.inpcrd) for each starting structure of the complex.

### Simulation and Refinement of the (DMA-135)-SLII complex structures

Initial (DMA-135)-SLII structures were minimized in sander using 2000 steps of steepest descent followed by 2000 steps of conjugate gradient. A 24 Å cutoff for non-bonded interactions was input alongside a 10 Å cutoff for calculation of the Born radii. After minimization, production MD simulations were performed using GPU-accelerated Particle Mesh Ewald Molecular Dynamics (pmemd) on NIVIDIA Tesla P100 GPUs. During this stage, NOE (20 kcal mol^−1^ Å^−1^) hydrogen-bonding and chirality restraints were used. Torsion angle restraints were manually incorporated to maintain the aromatic rings of DMA-135 in a planar orientation. The guanidinium group was assigned to its lowest-energy resonance structure. For the 50 ns simulation (total 25,000,000 steps of 2fs each), a non-bonded cutoff of 999.9 was used alongside a 10-Å cutoff for calculation of the Born radii. SHAKE was used to constrain bonds involving hydrogen and Langevin dynamics with a collision frequency of 2.0 ps^−1^ was used to control temperature. The RNA was heated from 0 to 300 K over 100ps and simulated for the remainder of the 50-ns simulation at 300 K.

After 50 ns of simulation, cpptraj was used to generate structures from the trajectory files, extracting the final frame of each simulation. (200 per starting structure). The 16 structures were simulated for 1ns in Amber tools 15 using sander and the ff99bsc0XOL3 force field. At this stage, both minimization and production used implicit solvent (igb=1), a salt concentration of 10mM, a non-bonded cutoff of 24 Å and a 10-Å cutoff for calculation of the Born radii. Residual Dipolar Couplings (RDCs) were incorporated into the refinement simulation with a weak-weighting coefficient of 0.01 as single-value restraints alongside the hydrogen bonding, NOE, and chirality restraints utilized in the previous phase of calculations. Structures were prepared once more in Antechamber and minimized over 4000 steps (2000 steps of steepest descent followed by 2000 steps of conjugate gradient). After this final minimization, the experimentally determined RDC values were fit to the structures by freezing the coordinates of the structure to allow generation of the molecular alignment tensor. In the simulation step, the structures were allowed to move in order to refine them according to available experimental restraints. Temperature scaling was arranged such that the system spent 1ns at 300K before cooling to 0K. Upon completion of this refinement step, the final frames from each of the 16 trajectories were extracted to individual *.pdb file. These final structures were ranked according to their total energies, with the lowest 10 represented here. Back-calculated RDCs were extracted from sander output files and assessed for their agreement with measured values. Once calculation completed, the 10 lowest energy structures for each construct were visualized in PyMOL and checked using MolProbity.

## Acknowledgements

The authors would like to acknowledge CCIC (Ohio State University) NMR facility manager Alexandar L. Hansen for assistance with setting up NMR experiments to measure RDCs and David Case (Rutgers University) for designing planarity restraints for DMA-135 in AMBER.

## AEH funding

Duke University; NIH U54 AI150470; NIH R35 GM124785 BST and GB funding: CWRU and Rutgers, NIH R01 GM126833

## Author contributions

N.N.P. synthesized all the amiloride derivatives described, designed and performed replicates of binding assays of SLII to Tat peptide and screening experiments with the amiloride derivatives.

Z.C. assisted in the design of and performed replicates of the binding assay of SLII to Tat peptide and screening experiments with the amiloride derivatives, and performed the titrations of small molecules using the Tat displacement assay.

J.D.C carried out biophysical (NMR and ITC) characterization studies of DMA-SLII and DMA-SLII-protein interactions and assisted with structural calculations of the SLII-(DMA-135) complex.

L.Y.C. assisted with biophysical characterization studies and performed structural calculations of the SLII-(DMA-135) complex.

A.S. performed structural calculations of the SLII-(DMA-135) complex and the AUF-SLII-(DMA-135) ternary complex.

S.P. completed NMR chemical shift assignments and structure calculations of AUF1.

M.L.L. helped conceived and completed all molecular and virology experiments and to analyze functional data.

A.E.H. helped conceive and plan experiments, particularly those related to small molecule synthesis and screening, and to analyze small molecule screening data.

G.B. helped conceive and plan experiments, particularly those related to the EV71 functional studies, and to analyze functional data.

B.S.T. helped conceive and plan experiments, particularly those related biophysical characterizations and structure calculations, and to analyze biophysical and NMR data.

B.S.T, A.E.H., G.B., M.L., N.N.P., J.D.C, L.Y.Y., and A.S. helped write the paper.

