## Supplemental Information for "Small Molecule Targeting IRES Domain Inhibits Enterovirus 71 Replication via an Allosteric Mechanism that Stabilizes a Ternary Complex"

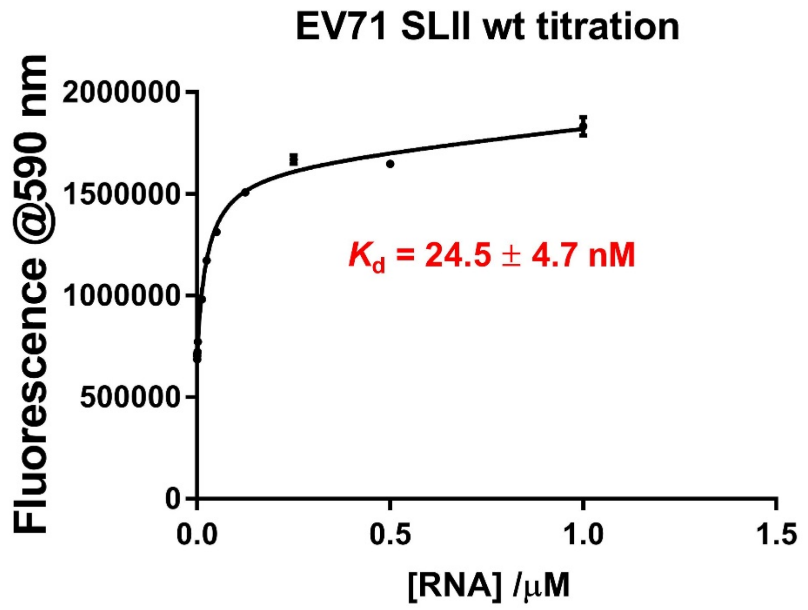

**Figure S1.** Binding assay for indicator peptide to SLII.

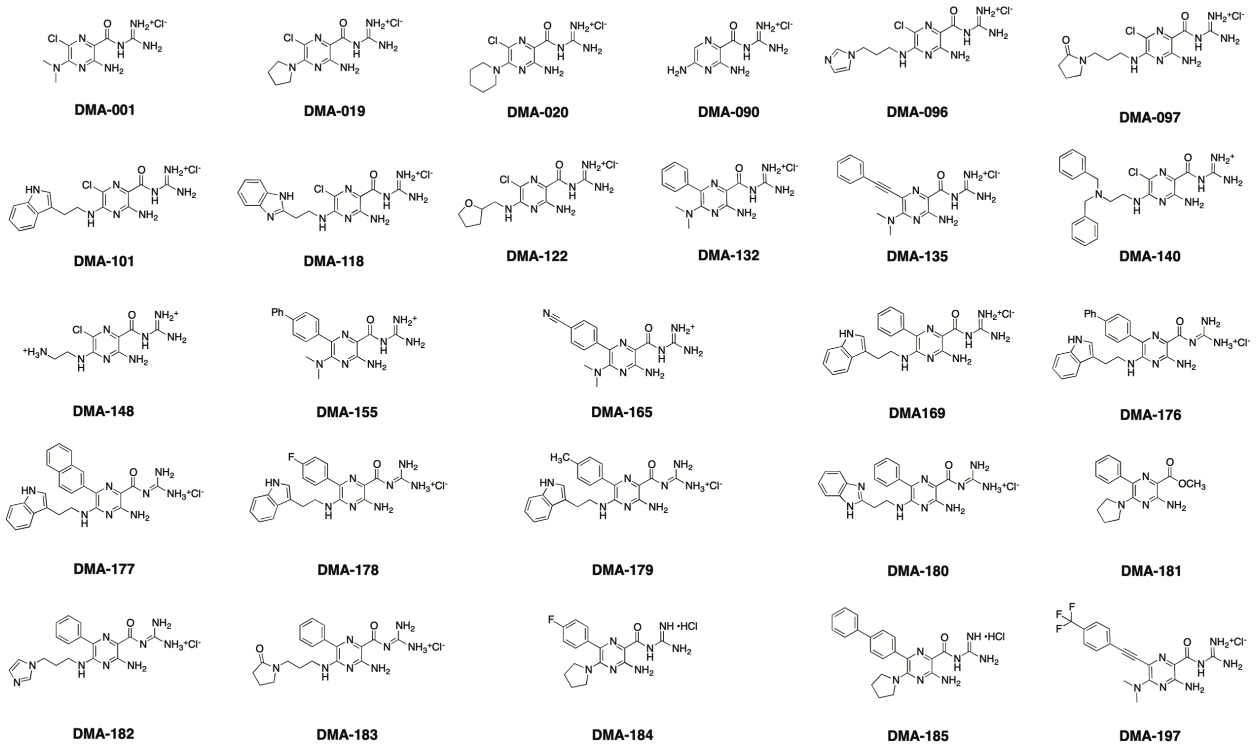

**Figure S2.** Structures of amiloride derivatives utilized in this study

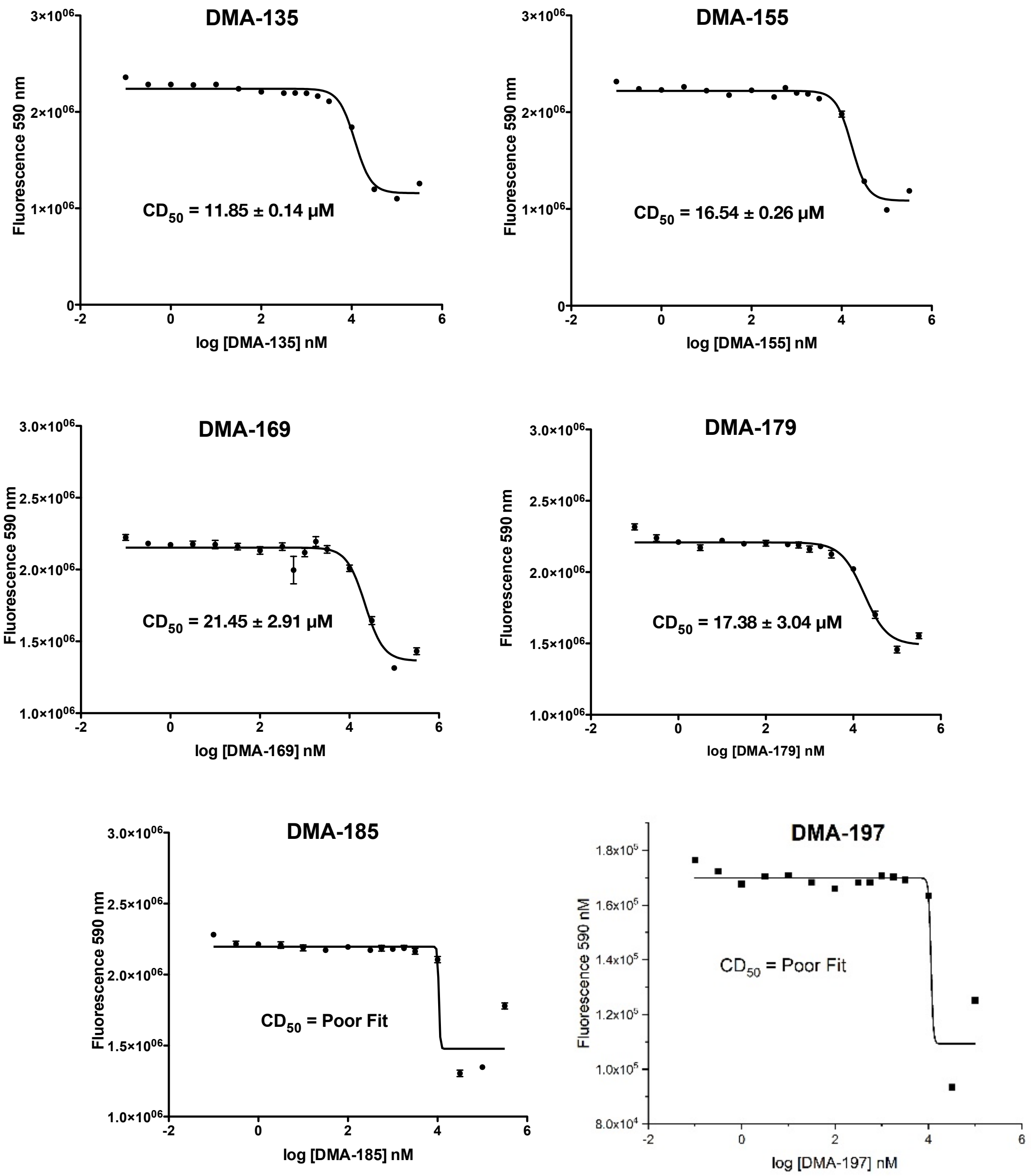

**Figure S3.** Full binding curves of amiloride molecules vs EV71 SLII RNA

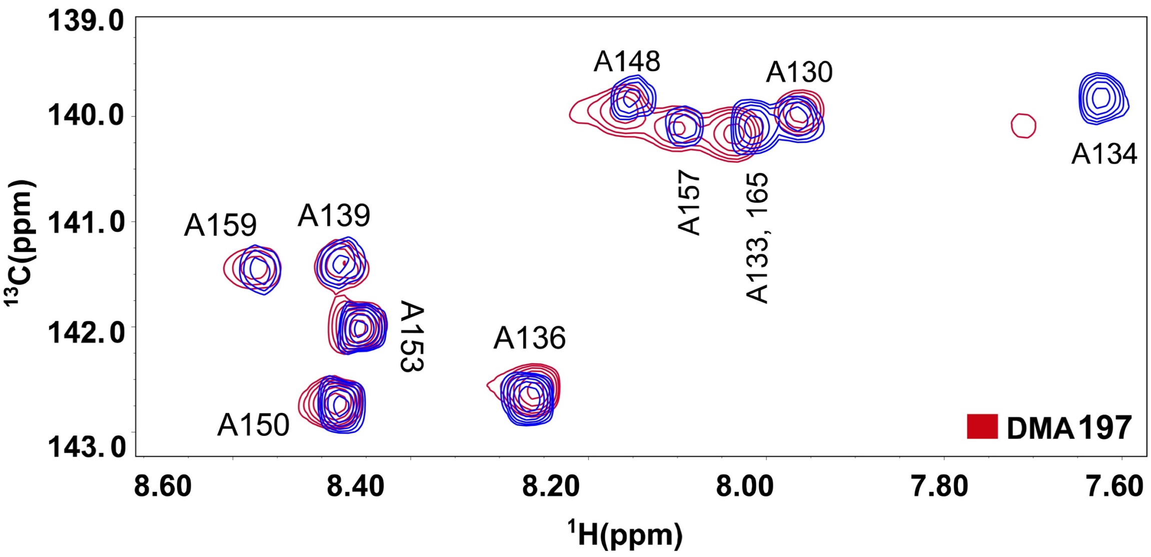

**Figure S4.** Single-point $^{1}$H-$^{13}$C TROSY HSQC titrations of A($^{13}$C)-selectively labeled SLII constructs with DMA-197 at 5-fold excess

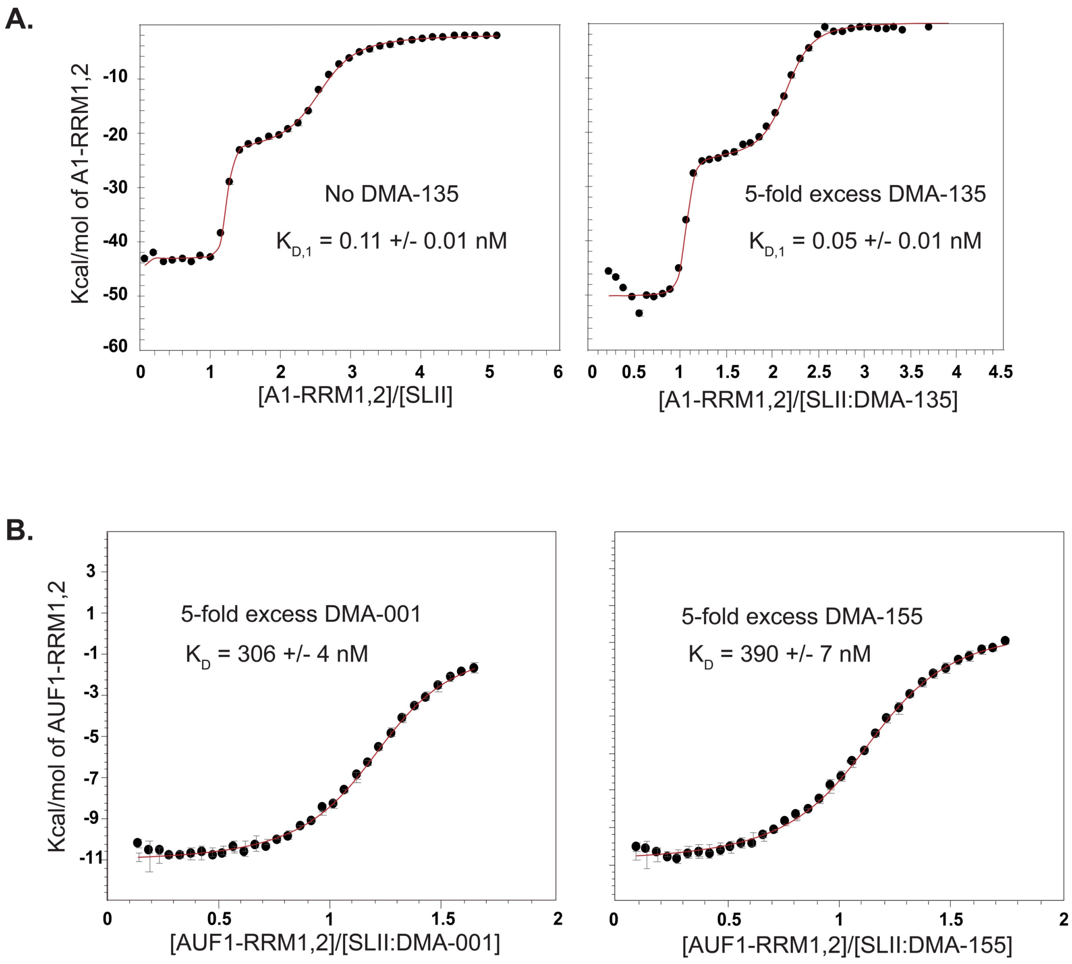

**Figure S5.** (A) Calorimetric titrations of A1-RRM1,2 into free SLII (left) and the SLII-(DMA-135) complex prepared at a 1:5 molar ratio. (B) Calorimetric titrations of AUF1-RRM1,2 into (left) SLII-(DMA-001) and (right) SLII-(DMA-155) each prepared at a 1:5 molar ratio. The experiments were performed in 10 mM $K_2HPO_4$, 20 mM KCl, 0.5 mM EDTA, and 4 mM DTT (pH 6.5) buffer at 298 K.

A.

B.

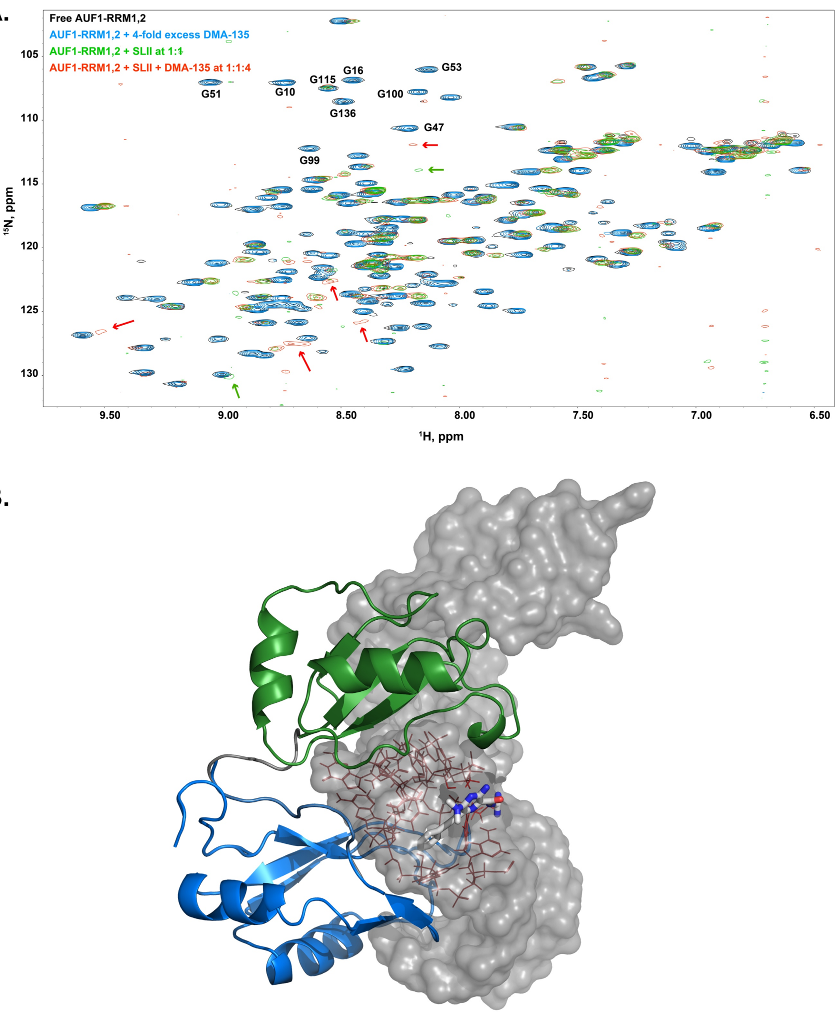

**Figure S6.** (A) Single-point $^{1}$H-$^{15}$N HSQC titrations of $^{15}$N-labeled AUF1-RRM1,2 (black) with excess DMA-135 (blue); with SLII added at a 1:1 molar ratio (green); or with the SLII-(DMA-135) complex prepared at a 1:4 molar ratio (red). Red arrows indicate new correlation peaks only observed in the presence of SLII-(DMA-135). (B) Structural model of the AUF1,2-SLII-(DMA-135) ternary complex. The structural model was calculated in HADDOCK using the observed AUF1-RRM1,2 chemical shift perturbations as docking restraints. The structure of AUF1-RRM1,2 is a preliminary solution NMR structure that will described elsewhere.

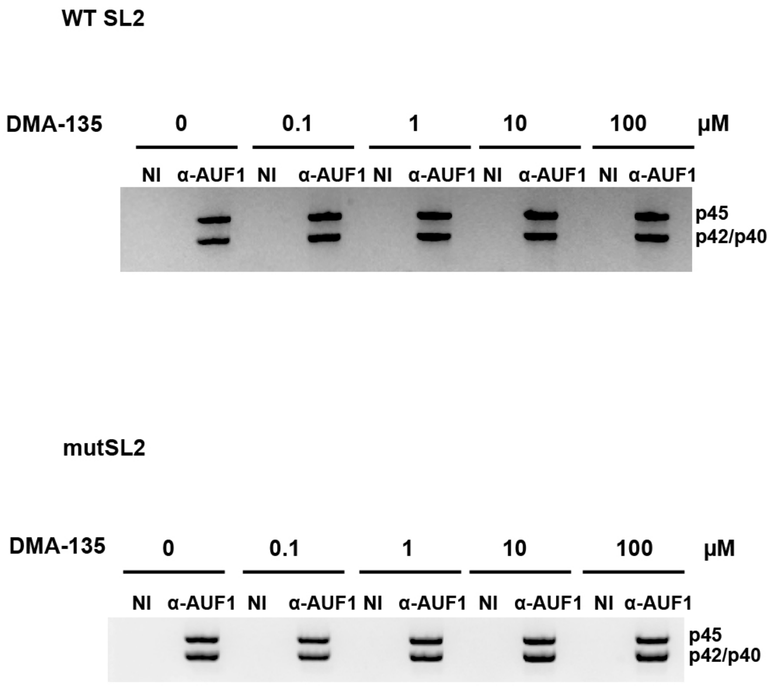

**Figure S7.** Aliquots of samples immunoprecipitated with either anti-AUF1 or non-immune serum (N.I.) from Figure 7C were analyzed by Western blot to verify anti-AUF1–dependent recovery of AUF1. The specific AUF1 isoforms are indicated. Note that immunoprecipitation using N.I. serum did not precipitate AUF1, as expected.

**Table S1.** $CD_{50}$ values for a selected set of small molecules

| **Molecule** | **$CD_{50}$ (μM)** |
|---|---|
| DMA-135 | 11.85 ± 0.14 |
| DMA-155 | 16.54 ± 0.26 |
| DMA-169 | 21.45 ± 2.91 |
| DMA-178 | 17.38 ± 3.04 |
| DMA-185 | Poor Fit |

**Table S2.** Summary of thermodynamic parameters for DMA-SLII and AUF1-SLII-(DMA-135) interactions.

| System | ΔG (kcal/mol) | ΔH (kcal/mol) | -TΔS (kcal/mol) | $K_D$ (nM) |
|---|---|---|---|---|
| **SLII to DMA-001** | -8.6 ± 0.4 | 12.8 ± 3 | -21.4 ± 2 | 584 ± 62 |
| **SLII to DMA-135** | -8.6 ± 0.1 | 12 ± 1 | -20.7 ± 0.4 | 525 ± 98 |
| **SLII to DMA-155** | -9.0 ± 0.1 | 11 ± 5 | -20 ± 5 | 271 ± 59 |
| **SLII to DMA-169** | No Data | | | |
| **SLII to DMA-178** | No Data | | | |
| **SLII to DMA-185** | No Data | | | |
| **AUF1 to SLII** | -8.8 ± 0.1 | -25.7 ± 0.7 | 16.8 ± 0.7 | 336 ± 44 |
| **AUF1 to SLII: DMA-135 (1:1.6)** | -9.0 ± 0.2 | -11.7 ± 0.5 | 2.7 ± 0.5 | 240 ± 72 |
| **AUF1 to SLII: DMA-135 (1:3.3)** | -9.34 ± 0.01 | -13.91 ± 0.04 | 4.58 ± 0.04 | 140 ± 2 |
| **AUF1 to SLII: DMA-135 (1:5)** | -9.37 ± 0.03 | -13.5 ± 0.2 | 4.1 ± 0.2 | 134 ± 7 |
| **AUF1 to SLII: DMA-001 (1:5)** | -8.88 ± 0.01 | -10.9 ± 0.4 | 2.1 ± 0.4 | 306 ± 4 |
| **AUF1 to SLII: DMA-155 (1:5)** | -8.73 ± 0.01 | -13.11 ± 0.01 | 4.38 ± 0.02 | 390 ± 7 |

**Table S3.** Summary of NMR-SAXS restraints and structure statistics

| | $SLII^{2231}$-DMA135 |
|---|---|
| Intra-residue | 162 |
| Sequential (\|*i*-*j*\|)=1 | 245 |
| Long-range | 24 |
| Hydrogen bonds | 75 |
| NOE restraints/residue | 12.34 |
| $C^H$ RDCs (base pair only) | 22 |
| Sugar Pucker | 155 |
| Amber Structure Statistics | |
| | $SLII^{2231}$-DMA135 |
| Average energy (cal $mol^{-1}$) | -9432.2872 |
| Overall RMSD | 3.7323 |
| RMSD lower and upper helices (hydrogen bond) | 2.4338 |
| RMSD lower helix | 0.2682 |
| RMSD upper helix | 0.3482 |
| NOE violations (> 0.4 Å) | 1.3 |
| Molprobity Clash Score (percentile) | 99-$100^{th}$ |
| PDB code | - |

*note: Amber structure statistics took 10 lowest structures to average.

**A. Characterization data for trifluoromethyl derivative DMA-197.**

**Amino(3-amino-5-(dimethylamino)-6-((4-fluorophenyl)ethynyl)pyrazine-2-carboxamido) methaniminium chloride (DMA-197)**: $^{1}$H NMR (500 MHz, Methanol-d4) δ 7.75 – 7.65 (m, 4H), 3.46 – 3.38 (m, 1H, NH), 3.37-3.32 (m, 6H); $^{13}$C NMR (126 MHz, MeOD) δ 165.5, 157.1, 155.6, 154.2, 131.3, 131.2, 126.7, 125.3, 112.8, 112.2, 90.2, 89.9, 39.4; HRMS (ESI+): Calculated for $C_{17}H_{17}F_3N_7O$ [M+H]: 392.1441, Found: 392.1448 (± 1.6 ppm). HPLC Analysis: Retention time = 7.106 min, Purity = 95.141%.

**$^{1}$H NMR Spectrum for Amino(3-amino-5-(dimethylamino)-6-((4-fluorophenyl)ethynyl)pyrazine-2-carboxamido) methaniminium chloride (DMA-197):**

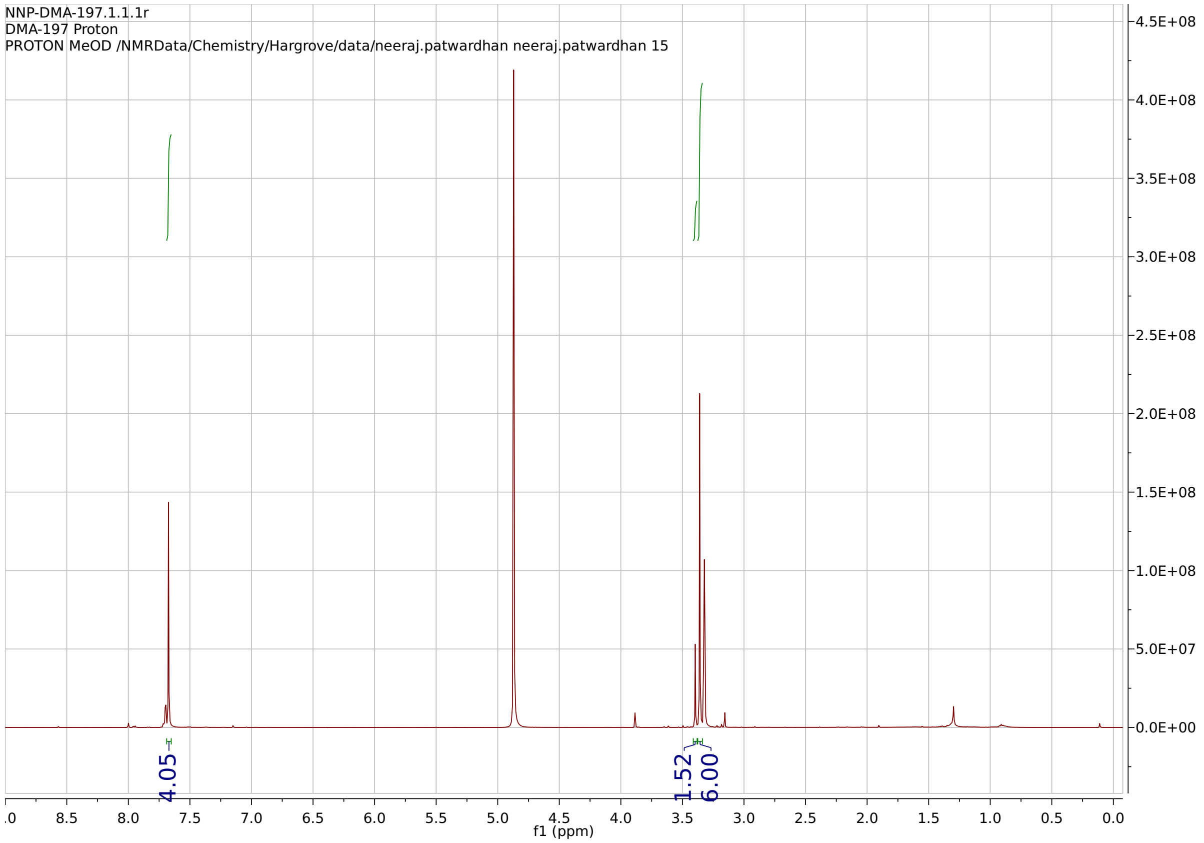

**$^{13}$C NMR Spectrum for Amino(3-amino-5-(dimethylamino)-6-((4-fluorophenyl)ethynyl)pyrazine-2-carboxamido) methaniminium chloride (DMA-197):**

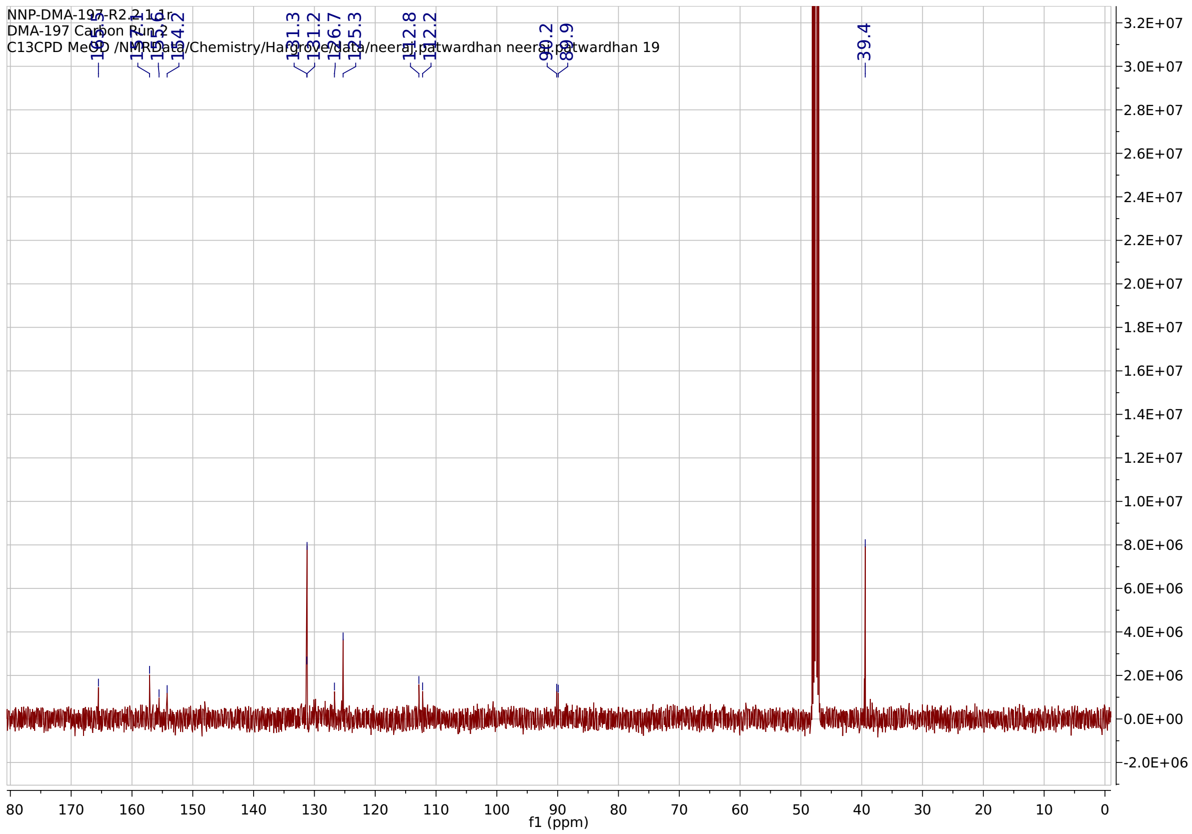

**HPLC Chromatogram for for Amino(3-amino-5-(dimethylamino)-6-((4-fluorophenyl)ethynyl)pyrazine-2-carboxamido) methaniminium chloride (DMA-197):**

### Analysis Report

#### <Sample Information>

| | | | |
|---|---|---|---|
| Sample Name | : NNPIII-129-R1 | | |
| Sample ID | : NNPIII-129-R1 | | |
| Data Filename | : NNPIII-129-R1.lcd | | |
| Method Filename | : NNP-Grd10-90_Slow_PDA_D2only.lcm | | |
| Batch Filename | : NNP_04_22_19_R1.lcb | | |
| Vial # | : 1-5 | Sample Type | : Unknown |
| Injection Volume | : 10 uL | | |
| Date Acquired | : 4/22/2019 11:02:53 AM | Acquired by | : chemist |
| Date Processed | : 4/22/2019 11:25:57 AM | Processed by | : chemist |

#### <Chromatogram>

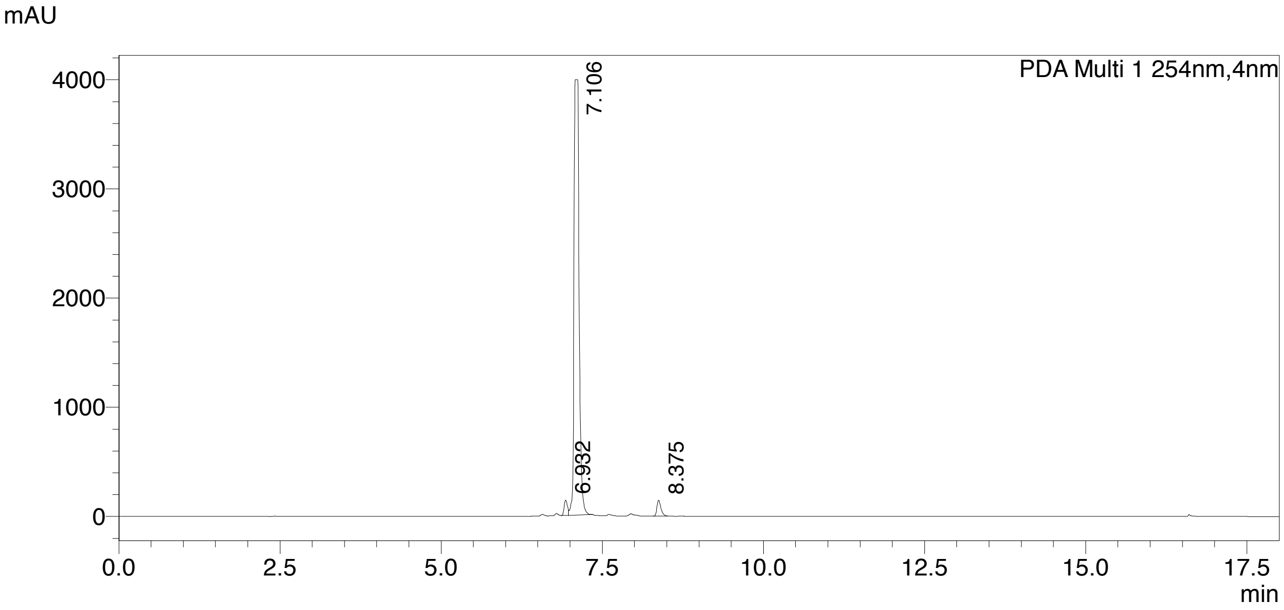

#### <Peak Table>

PDA Ch1 254nm

| Peak# | Ret. Time | Area | Area% |
|---|---|---|---|
| 1 | 6.932 | 497759 | 2.160 |
| 2 | 7.106 | 21925166 | 95.141 |
| 3 | 8.375 | 622086 | 2.699 |
| Total | | 23045010 | 100.000 |

#### B. Small molecule screening using the Tat peptide displacement assay –

**Z' Scores:** Z' Scores for each RNA: peptide system were calculated using the equation below, using 144 data points for each RNA:peptide complex –

$$Z' = 1 - \left( \frac{3\,(\sigma_{(positive)} + \sigma_{(negative)})}{|\,\mu_{(positive)} - \mu_{(negative)}|} \right)$$

$\mu_{(positive)}$ = Mean Fluorescence Intensity for peptide + RNA
$\mu_{(negative)}$ = Mean Fluorescence Intensity for peptide alone
$\sigma_{(positive)}$ = Standard deviation for positive
$\sigma_{(negative)}$ = Standard deviation for negative

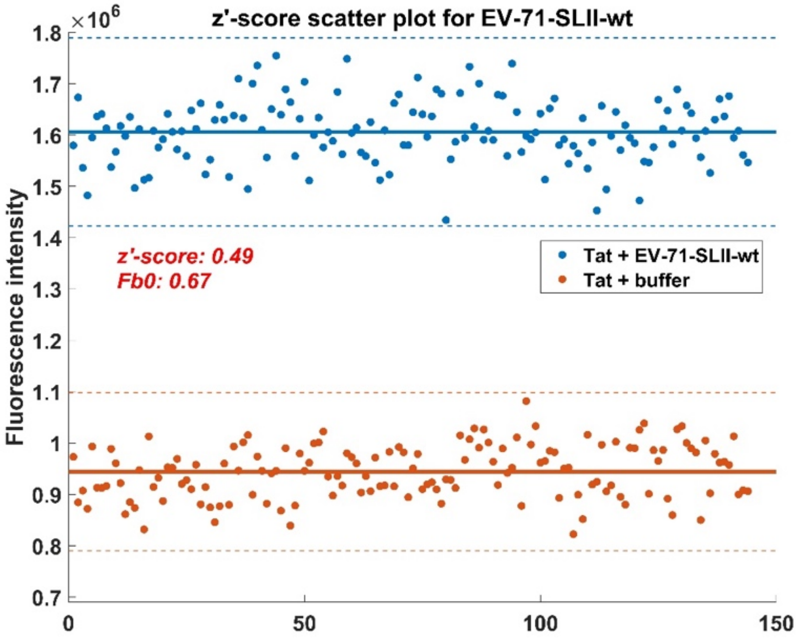

C. Screening results for panel of small molecules at 10 μM concentration:

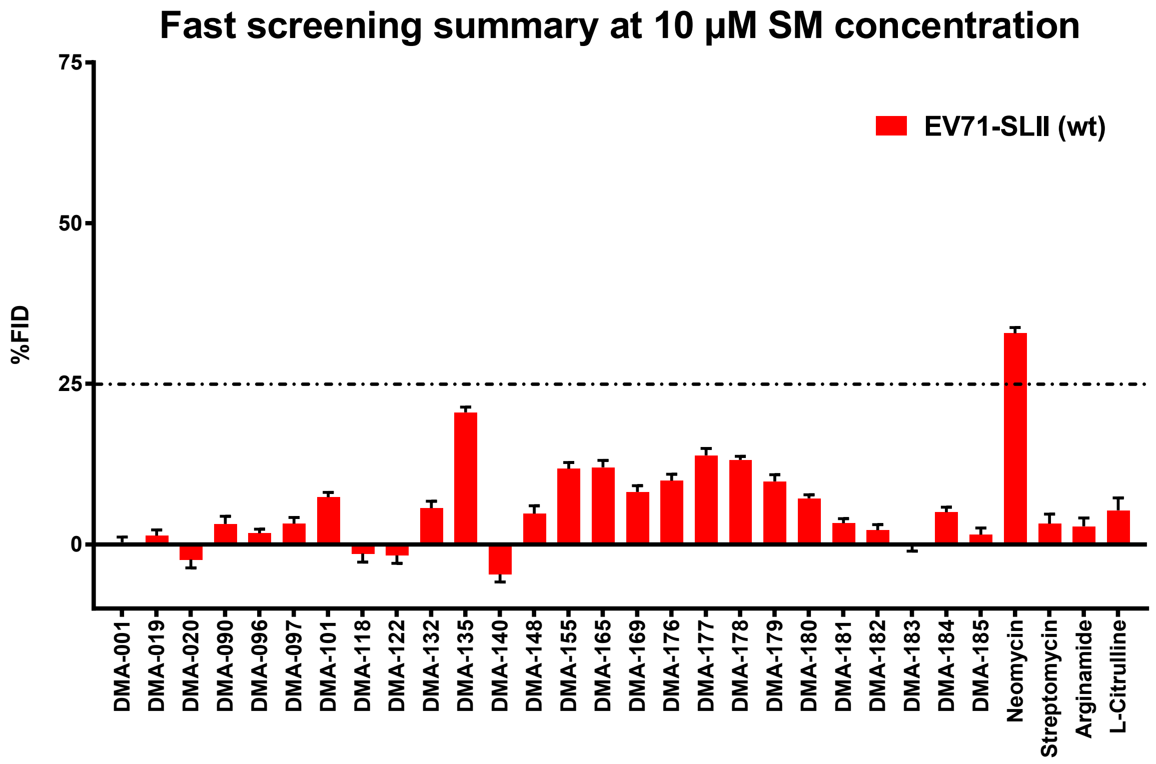

D. **Screening results for panel of small molecules at 50 µM concentration:**

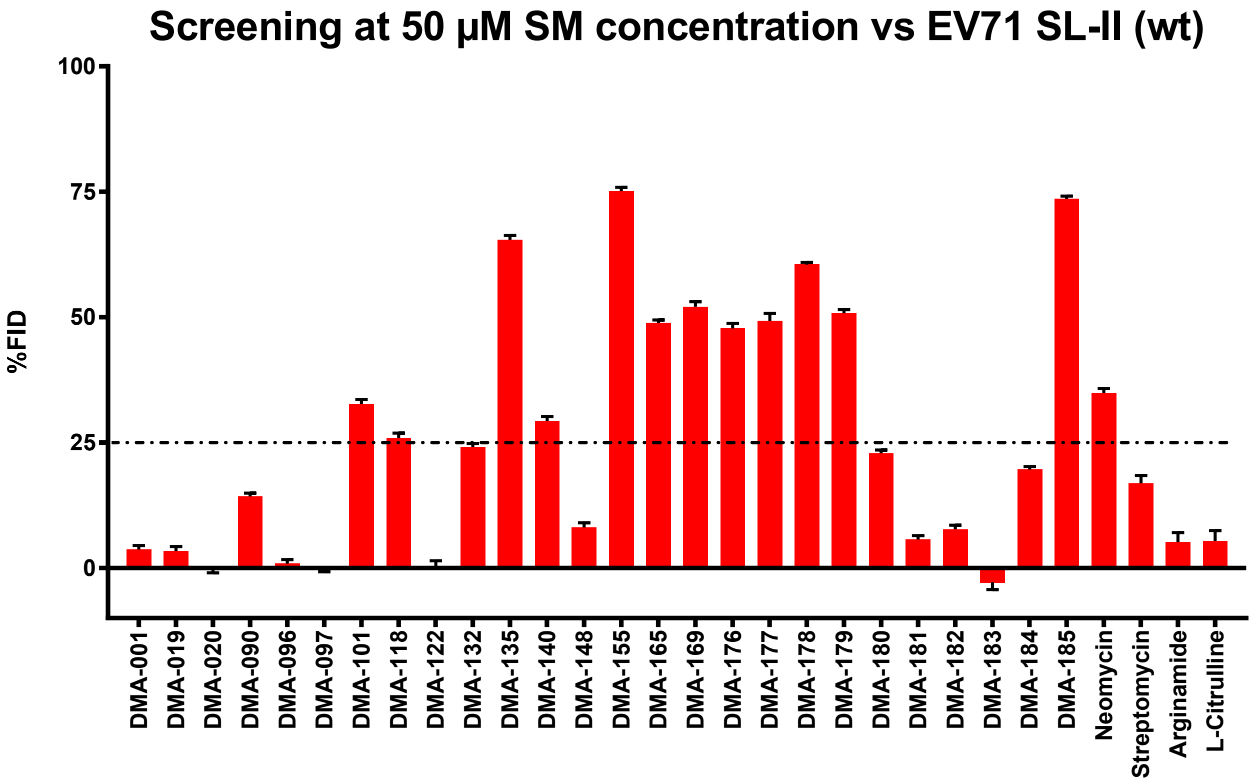
